# Enrichment of Extracellular Vesicles Subsets from Diverse Biological Sources Using Conventional and Image-Based Fluorescence Activated Sorting

**DOI:** 10.1101/2025.10.12.681862

**Authors:** Isabel Graf, Amanda Salviano-Silva, Jochen Behrends, Anne Rissiek, Christopher Urbschat, Santra Brenna, Hela Uplegger, Bente Siebels, Cecile L Maire, Kristoffer Riecken, Katrin Lamszus, Anke Diemert, Franz Ricklefs, Tim Magnus, Petra Arck, Berta Puig

## Abstract

Selective enrichment of extracellular vesicle (EV) subpopulations from the heterogeneous EV pool is essential for understanding their characteristic biological functions and exploiting their potential as diagnostic and prognostic biomarkers. However, isolation of specific EV-subsets remains challenging. Fluorescence-Activated Cell Sorting (FACS) has emerged as promising technique for EV subpopulations enrichment, despite limitations associated to their small size. Although FACS-based EV sorting has been reported, a broadly applicable and systematically validated workflow is still lacking.

Here, we describe and validate an optimized workflow for the sorting and analysis of EVs derived from diverse species, tissues, blood and cell culture systems. Using two advanced flow cytometric cell sorters, the BD FACSAria Fusion, and the BD FACSDiscover S8, we systematically evaluated key technical parameters, including nozzle size, sample dilutions, and sorting mode. The optimized workflow enabled efficient enrichment of differently labelled EV populations of interest, achieving near-100% purity, including rare subsets representing less than 10% of the total EV pool, while maintaining compatibility with downstream analyses. Sorted EV populations were characterized by high-sensitivity imaging flow cytometry, transmission electron microscopy, and liquid chromatography–tandem mass spectrometry. This workflow provides a robust framework for EV subset isolation and characterization, supporting both fundamental EV research and translational biomarker applications.

## INTRODUCTION

Extracellular vesicles (EVs) are secreted nanoparticles involved in cellular communication and are increasingly recognized as valuable biomarkers as well as promising tools for drug delivery. Because virtually all cell types release EVs, biological samples contain a highly diverse and heterogeneous EV population that differ in their cellular origins, molecular compositions, and biological function^1,2^. Studying EV cargo can provide important insights into the functional condition of the parent cells highlighting the potential of EVs as minimally invasive indicators for tissue analysis and disease monitoring through accessible biofluids^3,4^.

Developing methods capable of separating and analysing distinct EV subtypes is therefore crucial to acquire a deeper understanding of EV biology^5^. This is of particular relevance for clinical diagnostics and therapeutical applications, as only particular EV populations expressing defined markers or originating from particular cell types are of interest^5,6^. For example, the selective enrichment of tumour-derived EVs from the total EV pool in the plasma of brain cancer patients allows the detection of tumour-specific mutations, highlighting the importance of selective EV isolation for their use as liquid biopsies and biomarkers^4^. Yet, enriching specific EV-subpopulations remains challenging, especially when working with complex EV samples such as tissue or biofluids. Most of the standard methods used for EV isolation –such as differential centrifugation, density gradient ultracentrifugation, size-exclusion chromatography, or tangential flow separation- are based on separation by size and density and therefore cannot distinguish EV populations according to their cell of origin. While recent methodological advances in single-EV analysis including nanoscale flow cytometry or advanced imaging techniques such as Raman Tweezers Spectroscopy (RTS), Atomic Force Microscopy (AFM) or Cryo-Electron Microscopy (Cryo-EM) provide detailed characterization of EVs, these approaches do not enable the physical separation of EV subsets for subsequent untargeted cargo analysis^7^. Immunoaffinity-based approaches such as Magnetic Activated Cell Sorting (MACS) allows separation of distinct EV populations expressing predefined surface markers^8^. However, these methods provide limited flexibility for multiparametric discrimination and the simultaneous isolation of multiple EV subpopulations. In addition, residual magnetic beads or buffer-derived contaminants may be carried over during the isolation process and interfere with certain downstream analyses. These limitations may be particularly relevant when sample material is scarce and comprehensive characterization of multiple EV population is required.

Fluorescence-Activated Cell Sorting (FACS), also referred to as Fluorescence Activated Vesicle Sorting (FAVS) when applied to EVs, has emerged as a promising approach for the enrichment of specific EV subpopulations^9,10^. However, because conventional flow cytometric sorters (hereafter sorter) were originally designed for cell sorting, their application to EVs presents significant technical challenges, primarily due to the substantially smaller size and lower signal intensity of EVs compared to cells. Consequently, robust implementation of EV sorting requires careful optimization of instrument settings and experimental workflow.

Pioneering studies have successfully shown the feasibility of this approach^9–16^, however EV sorting is rarely performed, and it is still far from being standardized^17^. Reasons include limited reporting of necessary parameters, lack of proper validation, use of sorters that have since been discontinued or need of physical customisation of the flow cytometer. Moreover, while previous studies often demonstrated successful recovery of sorted EVs, assessment of enrichment in terms of purity has frequently been lacking. **Table 1** summarizes the technical parameters applied in previous studies and highlights the substantial heterogeneity in the experimental approaches and the limited degree of methodological standardization. Furthermore, how the EV source influences single-EV sorting has not been systematically investigated, particularly given the intrinsic heterogeneity of different sample types and the diversity of available labelling strategies. Therefore, there is an urgent need for a FACS-based EV-sorting workflow that is compatible with newer generation of sorters without requiring customization, facilitating straightforward implementation in core facilities with the in-house instruments and promoting broad applicability.

**Table 1.**
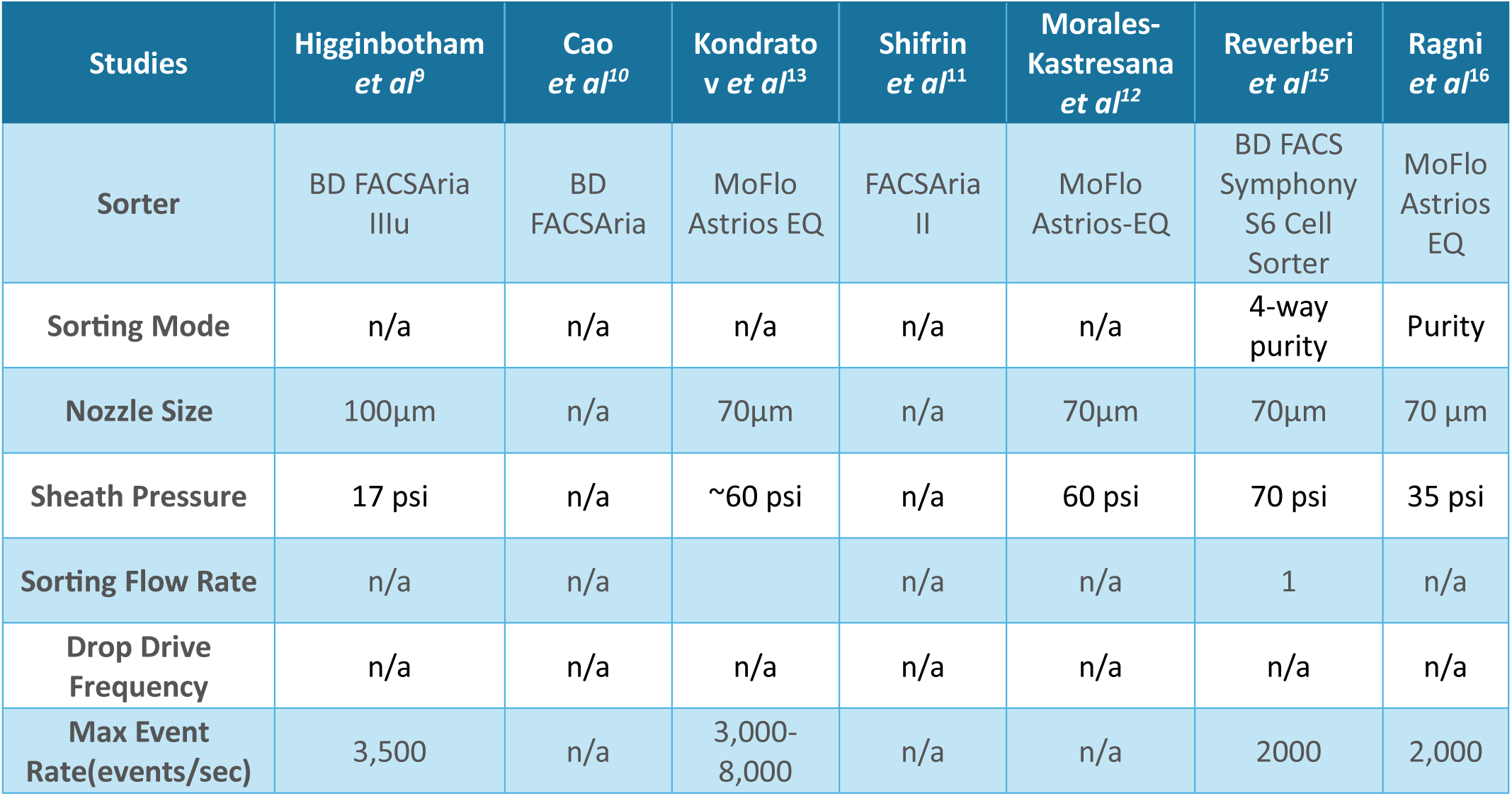
Overview of technical parameters reported for flow cytometric sorting of EVs in published studies.

In this study, we systematically evaluate and compare critical sorting parameters and establish a reproducible FACS-based workflow for a marker-based enrichment of defined heterogenous EV subsets using both, conventional and new-generation image-based spectral sorters. The goal of the present work is not to isolate EVs from biological sources but rather to establish a reproducible strategy for enriching specific EV subsets from pre-isolated EV preparations. We show the enrichment of EVs across a range of sources, including tissue-derived EVs, blood and cell culture conditioned media, as well as using different labelling strategies. These include labelling with membrane and intraluminal dyes, fluorescently labelled antibodies, and EVs endogenously expressing fluorescent reporter proteins, which may differentially influence the biophysical properties of EVs^18^. We demonstrate that the proposed workflow enables reliable enrichment of target EV populations across diverse sample types and labelling modalities while preserving vesicle integrity and compatibility with downstream analyses, including transmission electron microscopy (TEM) and proteomic profiling. Both the selective enrichment of EV subpopulations and their subsequent characterization by downstream analytical approaches are crucial for addressing fundamental questions in EV biology, including whether EV originating from distinct cell types carry unique cargo signatures and how these signatures change during disease progression^2,19–21^.

In summary, we present for the first time a robust framework for selective and reproducible enrichment of EV subpopulations across multiple biological sources and species, fluorescent labelling strategies, and FACS instruments. The workflow has strong potential for accelerating biomarker discovery in clinical settings and enabling in-depth characterization of EV subset functions in physiology and disease.

## METHODS

### EV isolation

#### Isolation of mouse tissue brain-derived EVs

Isolation of mouse tissue brain-derived EVs was performed following published methods^22^ with some modifications. Briefly, frozen mouse brain devoid of olfactory bulbs and cerebellum was gently cut with a scalpel, in RPMI-1640 medium (Gibco) containing 2mg/mL collagenase D (Roche) and 40U/mL of DNase I (Roche) on ice. Samples were then incubated at 37°C for 30 min with manual agitation every 5 min. Afterwards, protease inhibitors (Protease Inhibitors Cocktail, Roche) were added to stop the activity of the collagenase D.# The samples were homogenized by pipetting up and down for approximately 10 times with a 1,000 µL pipet with a cut tip and placed on a 70µm cell strainer (Fisher Scientific) on top of a Falcon tube to allow filtration by gravity, followed with a wash with 1mL of fresh RPMI. The filtrated samples were subjected to serial centrifugations at 4°C of 300xg for 10 min, 2,000xg for 20 min and 16,500xg for 20 min, always only collecting the supernatant for the next step with the exception of the last centrifugation where the resulting pellet was also collected, resuspended in 500µL of filtered (f)PBS containing protease inhibitors and kept at 4°C as “large/medium EVs”. The supernatants were placed in polypropylene centrifuge tubes (Beckman Coulter) and ultracentrifuged at 118,000xg for 150 min at 4°C (SW40Ti rotor, Optima Ultracentrifuge, Beckman Coulter). Afterwards, supernatants were discarded and the pellets containing “small EVs” were resuspended in 500µL of fPBS containing protease inhibitors. L/m EVs and sEVs preparations were mixed (final volume of 1mL), mixed with Optiprep (Axis Shield) to reach a concentration of 45%, loaded at the bottom of an ultracentrifuge propylene tube and layers of 30% and 10% of Optiprep were layered on top. Samples were then centrifuged at 186,000xg for 150 min at 4°C as described above and the white band at the interface between 10% and 30% of the cushion gradient was collected as the isolated EV fraction (2 mL). This fraction was further diluted in cold fPBS and centrifuged again at 118,000xg for 150 min at 4°C to finally collect a pellet enriched in tissue brain-derived EVs. This pellet was resuspended in 100µL of fPBS containing protease inhibitors and kept at -80°C in aliquots for further experiments.

All animal experiments were approved by the local animal care committee (Behörde für Justiz und Verbraucherschutz of the Freie und Hansestadt Hamburg, project number N105/2023 and ORG1055) and in compliance with the guidelines of the animal facility of the University Medical Center Hamburg-Eppendorf.

#### Isolation of human blood-circulating EVs

Venous blood samples of healthy pregnant individuals (3^rd^ trimester) and non-pregnant individuals were obtained via peripheral venipuncture with a butterfly needle (21 gauge) collection system. Samples were rested for minimum 60 min to allow RBC clotting. The RBC clot was subsequently pelleted by centrifugation at 2,000 g for 15 min and serum was aspirated. Serum samples were stored at −80 °C in 250µl aliquots and were not thawed before usage. Detailed information on preanalytical values, blood collection and processing as well as quality controls are reported according to the ‘MIBlood-EV. Standardized Reporting Tool for Blood EV Research’ as suggested by the Guidelines of the International Society of Extracellular Vesicles (**Supplementary Table 1**)^23,24^.

Next, to pellet cell fragments, larger particles and other debris, the serum samples were diluted 1:2 with fPBS and centrifuged at 10,000xg for 30 min. The supernatant was transferred into an 8,9 mL Beckmann Coulter ultracentrifugation tube and filled up with fPBS for ultracentrifugation at 100,00x g for 70 min using a Type 70.1 Ti Fixed-Angle Titanium Rotor. Subsequently the pelleted medium and small EVs were resuspended in fPBS for further processing.

Samples of healthy pregnant individuals were obtained from the PRINCE study (Prenatal Identification of Children’s Health), an ongoing prospective longitudinal pregnancy cohort study. Healthy non-pregnant individuals were recruited at the University Medical Center Hamburg-Eppendorf, Germany. All study subjects signed informed consent forms. The PRINCE study protocol was approved by the ethics committee of the Hamburg Chamber of Physicians under the license number PV 3694 and was conducted according to the Declaration of Helsinki for Medical Research involving Human Subjects.

#### Isolation of human and mouse glioma-derived EVs

The human glioma cell line NCH1681 (IDH1^R132H/WT^)^25^ and the mouse glioma line Mut3 (hGFAP-cre; Nf1flox/+; Trp53−/+)^26^ were used. NCH1681 was transduced with lentivirus (CSCW2 lentiviral backbone) encoding for palm-tdTomato, while Mut3 cells were transduced with mTagBFP2. Cells not expressing tdTomato or BFP2 were removed by flow cytometry using a BD Aria Fusion cell sorter. Cells were grown for up to 10 days in serum-free Neurobasal Medium (NBM) with B-27 Supplement, FGF-2, EGF and Heparin.

Conditioned media was collected and centrifuged at 300xg for 7 min. The cell-depleted supernatant was cleared from remaining debris by centrifuging at 2,000xg for 10 min, and then followed by ultracentrifugation at 100,000xg for 70 min at 4°C to isolate small/medium EVs. EV pellets were carefully resuspended in fPBS and maintained at 4°C during the whole isolation protocol.

### Validation of EV isolation

All isolated EVs were characterized according to the most recent guidelines^24^. Concentration and size distribution were evaluated using Nanoparticle Tracking Analysis (NTA), morphology by TEM, and the presence of classical EV markers (CD9, CD81 and CD63) was assessed by Imaging Flow Cytometry (IFCM).

#### NTA

The concentration and size of EVs before and after sorting were determined by NTA, using a LM10 instrument (Nanosight, Amesbury, UK). Before sorting conditions: brain tissue-derived EVs were diluted in fPBS (1:1000) and ten movies of 0.5 min each were recorded on camera level 16 and analyzed with detection threshold 6 and screen gain 2. Human blood-circulating EVs were diluted in fPBS (1:200) and five movies of 1 min each were recorded on camera level 14 and analyzed with detection threshold 5 and screen gain 2. Human and mouse glioma-derived EVs were diluted in fPBS (1:300), and five movies of 1 min each were recorded on camera level 15 and analyzed with detection threshold 4 and screen gain 2.

After sorting conditions: NTA measurements of each sample type were recorded with the same settings without any dilutions.

The analysis was performed by NTA 3.0 software. Particle concentrations were normalized to particles per mL, and particle size is reported as modal diameter.

#### Negative Staining and TEM

A volume of 30 µl of sorted EVs was diluted with 16% paraformaldehyde (PFA) in PBS to achieve a final concentration of 4% PFA. 5 µl of this fixed EV solution was applied to a Formvar-carbon coated 200 mesh copper grid (#ECF200-Cu-50) (previously treated with GLOQUBE PLUS to increase the positive charge). and allowed to rest in a dry environment for 20 min. The samples were then rinsed in PBS three times for two min and subsequently stained with a 2% methylcellulose-uranyl acetate solution for 10 min on ice. The excess solution was then removed by gently looping the grid onto a filter paper. EVs were analyzed and imaged using the Jeol JEM2100Plus TEM, equipped with a XAROSA CMOS camera.

#### Fluorescent sample labelling for IFCM and EV sorting

##### Mouse tissue brain-derived EVs

For IFCM, EVs were incubated for 30 min in the dark at 4°C with normal rat serum and anti-mouse CD16/32 (TruStain fcX, BioLegend) to block unspecific FcγRII/III binding. The antibody cocktail included APC anti-CD9 (BioLegend, clone MZ3, 0.2 mg/mL) diluted 1:10, APC anti-CD63(BioLegend, clone NVG-2, 0.2 mg/mL) and APC anti-CD81 (BioLegend, clone Eat-2, 0.2 mg/mL) anti-mouse antibodies. The antibody cocktail was filtered using 300 kDa filter (Nanosep) at 7,000xg for 10 min at 4°C and EVs were stained for 45 min in the dark at 4°C. Subsequently the EVs were washed using the 300 kDa filter mentioned before at 7,000xg for 10 min at 4°C and resuspended in fPBS.

For the sort, EVs were stained with mCling-Atto 647, a dye that is retained at the plasma membrane and endocytic membranes^27^ and with CFSE, a cell permeable dye that binds covalently to intracellular proteins. For mCling labelling, EVs were incubated with mCling (Synaptic Systems, final conc. 0.4µM) on ice for 5 min in the dark. The labelling reaction was stopped by adding 1% BSA. For CFSE (Thermo Fisher Scientific, final conc. 0.5µM) labelling, EVs were incubated for 2 h at 37 °C in the dark. Following dye incubation, the EV suspensions were transferred to ultracentrifugation tubes containing 11 mL fPBS and ultracentrifuged at 118,000xg for 150 min at 4 °C. The EV pellets were subsequently resuspended in fPBS.

##### Human blood-circulating EVs

For IFCM, EVs were treated with human IgG (ChromPure, Jackson ImmunoResearch) to block unspecific FcγRII/III binding for 30 min in the dark at 4°C diluted in 8% exosome depleted fetal calf serum (FCS, gilbeco) in fPBS. The antibody cocktail included Pacific Blue anti-CD9 (BioRad, clone MM2/57, 0.05 mg/mL) diluted 1:10, Pacific Blue anti-CD63 antibody (BioLegend, clone H5C6, 200µg/mL), Pacific Blue anti-CD81 antibody (BioLegend, clone TAPA-1, 300µg/mL) anti-human antibodies. Again here, antibody cocktail was filtered using 300 kDa filter (Nanosep) at 7,000xg for 10 min at 4°C and subsequently the EVs were stained for 45 min in the dark at 4°C. After the incubation EVs were washed using the 300 kDa filter mentioned before at 7,000xg for 10 min at 4°C to remove unbound antibodies and resuspended in fPBS. This sorted population was then assessed by IFCM using the same protocol with the following anti-human antibodies: FITC anti-CD41 (Biolegend, clone HIP8, 20µg/mL), FITC anti-ApoB (BioLegend, clone A-6, 200 µg/mL) and PE anti-CD9 (BioLegend, clone H19a, 20 µg/mL).

For sorting, the EVs same staining procedure was applied, except that the FCS was omitted, to prevent high albumin levels in the downstream mass spectrometric analysis. The following anti-human antibodies were used for the sort: AF647 anti-CD34 (Bioss, 1µg/µl, clone 4H11) diluted 1:10 and PE anti-CD9 (Biolegend, H19a clone, 20µg/mL) diluted 1:10.

##### Human and mouse glioma-derived EVs

For IFCM, human-EVs samples (positive for palm-tdTomato; acquired in Channel 03), were stained with a cocktail of the following anti-human antibodies conjugated with FITC (3μL of each): anti-CD9 (BioLegend, clone HI9a, 20μg/mL) diluted 1:30, anti-CD81 (BioLegend, clone 5A6, 200μg/mL), and anti-CD63 (BioLegend, clone H5C6, 200μg/mL), in 3μL of fPBS containing 8% exosome-depleted FBS (Invitrogen, cat. no. A2720801), for 45 min at room temperature (RT) in the dark. Previous to the incubation, the antibody cocktail was filtered using 300 kDa filter (Nanosep) at 7,000xg for 10 min at 4°C to eliminate aggregates. The same procedure was used for glioma-derived murine samples (positive for BFP2; acquired in Channel 07), but using the following anti-mouse antibodies conjugated with AlexaFluor 647: anti-CD9 (BioLegend, clone MZ3, 0.5mg/mL), anti-CD81 (BioLegend, clone Eat2, 0.5mg/mL), and anti-CD63 (BioLegend, clone NVG-2, 0.5mg/mL).

Human-derived EVs (positive for palm-tdTomato) and murine-derived EVs (positive for BFP2) were sorted without any additional staining.

### Imaging Flow Cytometry (IFCM) – technical settings

For IFCM we used the Cytek Amnis ImageStream Mk II. All samples were measured at 60x magnification at low flow rate. The following channels were used for acquisition: FITC and CFSE – channel 02, PE and palm-tdTomato – channel 03, BFP2 – channel 07, mCling and AF647 – channel 11. All samples were analyzed using IDEAS software version 6.2 (Amnis, Luminex Corporation).

### Step-by-step setup for EV sorting with the BD FACSAria Fusion and BD FACSDiscover S8

#### 1 Instrument-setup

- BD FACSAria Fusion:

i. Clean the flow cell with Contrad 70 and incubate for approximately 15 min. Thoroughly flush the system by running the stream at a high flow rate before loading FACS Clean and incubating for an additional 15 min. Perform a final rinse with distilled water at a high flow rate. Center the stream of sorted droplets relative to the collection tube to minimize the risk of sorted droplets adhering to the tube wall.
ii. Center the stream of sorted droplets relative to the collection tube to minimize the risk of sorted droplets adhering to the tube wall.
iii. Set collection chamber and sample tube holder temperature to 4°C.
- BD FACSDicover S8:

i. Perform startup procedure twice, first run the extended fluidics startup, second run the daily fluidics startup procedure.
ii. Run the sample line backflush twice, to minimize background.
iii. Set collection chamber and sample tube holder temperature to 4°C.

#### 2 Instrument calibration

- BD FACSAria Fusion:

i. Calibration beads (BD CS&T Beads) to optimize the time delay, essential for sorting small particles.
ii. 7.5µm BD FACS Accudrop Beads to optimize the drop delay, repeating as often as necessary until the drop delay stabilizes.
- BD FACSDicover S8:

i. Calibration beads (BD FACSDiscover Setup Beads) enable the software to automatically characterize, track, and report performance measurements in conjunction with onboard LEDs.
ii. BD Cell View Calibration Beads enable the software to automatically calibrate the imaging system.
iii. 7.5µm BD FACS Accudrop Beads to optimize the drop delay.

#### 3 Setup parameters for sorting

- For nozzle size, sheath pressure, sorting flow rate and event rates, please refer to **Table 2**.
- Sorting mode mask: Purity^28^: Up to two consecutive drops may be sorted, provided that the following drops are free of contamination. This mode prioritizes sample purity at the expense of yield.
- Thresholds used at the BD FACSAria Fusion :

i. Human and mouse glioma-derived EVs: BV421 OR PE
ii. Human blood-derived EVs: PE OR APC
iii. WT mouse brain-derived EVs: APC OR FITC

OR logic: one of the specified parameters exceeds the threshold

**Table 2.** Technical features and physical requirements enabling successful EV sorting.

| Feature | Physical requirement | Ref | Possible settings | Selected settings |
| --- | --- | --- | --- | --- |
| **Sample dilution**<br>(maximum event rate/min) | High dilutions decrease the probability of having more than one EV at the interrogation point, allowing single EV analysis and avoiding coincidence (swarming effect). | 17,35 | ∞ | **Fusion:** 10,000<br>**S8:** 2,000 |
| **Core stream & sorting flow rate** | Minimum possible diameter to ensure individual EVs pass one at a time. through the laser beam. The core stream diameter depends on the pressure difference between the sheath fluid -the outer fluid that surrounds the sample stream- and sample fluid -the inner stream containing the EVs-. By applying a low flow rate this difference can be maintained small, producing a relatively narrow stream. | 28 | **Fusion:**<br>1-11<br>**S8:**<br>1-50 | **Fusion:** 1<br>**S8:** 1 |
| **Nozzle size, sheath pressure and drop drive frequency** | The smaller the nozzle, the higher the pressure, resulting in faster droplet generation rate and smaller droplets.<br>Trade-off between small droplet size, increasing the chance for single EV sorting, and high pressure, which might impact the EVs. High shear forces in cell sorting have been shown to impact cells. | 14,24 | **Nozzle size:**<br>**Fusion** [μm]<br>70, 85, 100, 130<br>**S8** [μm]:<br>85, 100, 130 | **Fusion:**<br>Nozzle: 100μm,<br>Sheath pressure: 20 psi,<br>Drop drive freq: 30Khz<br>**S8:**<br>Nozzle: 100μm,<br>Sheath pressure: 20 psi<br>Drop drive freq: 34Khz |
| **Sort precision mode** | Sort modes (masks) provide different combinations of particle deflection settings such as yield and purity.<br>Yield/Enrichment defines how close to the edge of the drop, a particle of interest can be located before sorting an additional drop<br>Purity: defines how close, a contaminating drop can be located before ignoring the drop being interrogated | 28 | **Fusion:**<br>Purity, 4-way purity, yield, single cell<br>**S8:**<br>Purity enrichment, single cell | **Fusion & S8: Purity**<br>Purity is set to maximum: only drops free of contamination (within the purity mask) are sorted, resulting in a sorted sample of high purity, at the expense of recovery and yield. |

All thresholds were set at 200.

- Thresholds used at the BD FACSDicover S8:

i. Human and mouse glioma-derived EVs: Threshold SSC (Imaging), 1,009
ii. Human blood-derived EVs: Threshold SSC (Imaging), 5,075
iii. WT mouse brain-derived EVs: Threshold SSC (Imaging), 7,110

#### 4 Preparing the instruments for measurement

- BD FACSAria Fusion: Clean the flow cell with Contrad 70 and incubate for approximately 15 min. Thoroughly flush the system by running the stream at a high flow rate before loading FACS Clean and incubating for an additional 15 min. Perform a final rinse with distilled water at a high flow rate.
- BD FACSDicover S8: Two times backflush; run a sample with rinse for 15 min with highest flow rate; two times backflush, run a sample with water for 15 min with highest flow rate; reduce flow rate to 1 and run the water sample for 10 min; two times backflush.

*Both BD FACSAria Fusion and* BD FACSDicover *S8:*

Between the samples, backflush twice to avoid contamination from previous samples, followed by running 1 min rinse and 1 min water, followed by a final backflush.

#### 5 Set-up of gates including controls

A detailed description of the gating strategy can be found in the main manuscript.

#### 6 Sample measurement

To stabilize the stream/pressure after cleaning procedures, the samples were measured at least 1 min on the instruments before recording the data. Data was collected for 1 min with the flow rate of 1 (for both instruments).

### Proteomic Analysis using Liquid Chromatography Tandem Mass Spectrometry (LC-MS/MS)

Samples were dissolved in lysis buffer containing 100 mM triethyl ammonium bicarbonate (TEAB) and 1% w/v sodium deoxycholate (SDC) buffer, boiled at 95 °C for 5 min and sonicated. Disulfide bonds were reduced in 10 mM dithiothreitol for 30 min at 56 °C and alkylated in presence of 20 mM iodoacetamide for 30 min at 37 °C in the dark. Then, the samples were dissolved to a concentration of 70% acetonitrile (ACN) and carboxylate modified magnetic E3 and E7 speed beads (Cytvia Sera-Mag™, Marlborough, USA) at 1:1 ratio in LC-MS grade water were added in a 10:1 (beads/protein) ratio adapted from the SP3-protocol workflow^29^. Samples were shaken at 1,400 rpm for 18 min at RT. Tubes were placed on a magnetic rack and the supernatant was removed. Magnetic beads were washed two times with 100% ACN and two times with 70% ethanol on the magnetic rack. After resuspension in 50 mM ammonium bicarbonate, digestion with 100 ng trypsin was performed (sequencing grade, Promega) at 37 °C overnight while shaking at 1,400 rpm. Tryptic peptides were bound to the beads by adding 95% ACN and shaken at 1,400 rpm for 10 min at RT. Tubes were placed on the magnetic rack, the supernatant was removed, and the beads were washed two times with 100% ACN. Elution was performed with 2% DMSO in 1% formic acid (FA). The supernatant was dried in a vacuum centrifuge and stored at -20°C until further use.

Chromatographic separation of peptides was achieved with a two-buffer system (buffer A: 0.1% FA in H_2_O, buffer B: 0.1% FA in 80% ACN) on a UHPLC device (VanquishTM neo UHPLC system, Thermo Fisher). Attached to the UHPLC was a peptide trap cartridge (300 µm x 5 mm, C18, PepMap™ Neo Trap Cartridge, Thermo Fisher) for online desalting and purification, followed by a 25 cm C18 reversed-phase column (75 µm x 250 mm, 120 Å pore size, 1.7 µm particle size, Aurora Ultimate, IonOptics). Peptides were separated at a flow rate of 0.4 µL/min using a 70 min method with linearly increasing ACN concentration from 3% to 34% ACN over 60 min.

MS/MS measurements were performed on a quadrupole-orbitrap hybrid mass spectrometer (Exploris 480, Thermo Fisher Scientific). Eluting peptides were ionized using a nano-electrospray ionization source (nano-ESI) with a spray voltage of 1,800 and analysed in data independent acquisition (DIA) mode. For each MS1 scan, ions were accumulated until 3 x 10^6^ ions (AGC Target) was reached in automatic mode. Fourier-transformation based mass analysis of the data from the orbitrap mass analyzer was performed covering a mass range of m/z 400 – 800 with a resolution of 120,000 at m/z 200. Within a precursor mass range of m/z 400-800 fragmentation in DIA-mode with m/z 12 isolation windows and m/z 1 window overlaps were performed. Fragmentation was performed at normalized collision energy of 30% using higher energy collisional dissociation (HCD). Orbitrap resolution was set to 60,000 with a first mass of m/z 120.

LC-MS/MS data were searched with the CHIMERYS DIA algorithm integrated into the Proteome Discoverer software (v3.1.0.638, Thermo Fisher Scientific) against a reviewed human or murine Swissprot database using Inferys 3.0 fragmentation as prediction model. Carbamidomethylation was set as a fixed modification for cysteine residues. The oxidation of methionine was allowed as variable modification. A maximum number of one missing tryptic cleavage was set. Peptides between 7 and 30 amino acids were considered. A strict cutoff (FDR< 0.01) was set for peptide identification. Quantification was performed by CHIMERYS based on fragment ions.

Gene ontology analysis was performed using the STRING database^30^.

The mass spectrometry proteomics data have been deposited to the ProteomeXchange Consortium via the PRIDE^31^ partner repository with the dataset identifier PXD069156.

## RESULTS

The key objective of this study was to determine the technical parameters that enable selective sorting of EVs using flow cytometric sorting platforms. To systematically evaluate critical instrument settings and technical parameters, we selected the most complex sample type evaluated within this study, i.e., EVs derived from brain tissue. These samples exhibit substantial heterogeneity and background complexity, thus providing a stringent test condition for workflow optimisation. Once optimal conditions were established, the same parameters were applied to other EV sources, including circulating blood–derived EVs and cell culture–derived EVs to validate the robustness and transferability of the workflow across different EV types and sources. The experimental workflow to establish the optimal EV sorting conditions is illustrated in **Figure 1**. Prior to and after sorting, EV samples were characterized with IFCM to assess the degree of purity and EV recovery relative to the input samples. IFCM is a highly sensitive flow cytometry-based imaging technique for single EV analysis^32,33^. A major advantage of IFCM is that false-positive fluorescent events, which may display a sideward-scatter and fluorescence profile similar to those of EVs, can be readily identified as they can be distinguished through the corresponding brightfield images (**Supplementary Figure 1A-C**). This enabled verification that the sorted particles were bona fide EVs. The IFCM gating strategy applied within our experimental set-up to exclude doublets, swarmed EVs and false-positive particles with inadequate brightfield image, is depicted in **Supplementary Figure 1D.**

**Figure 1.**
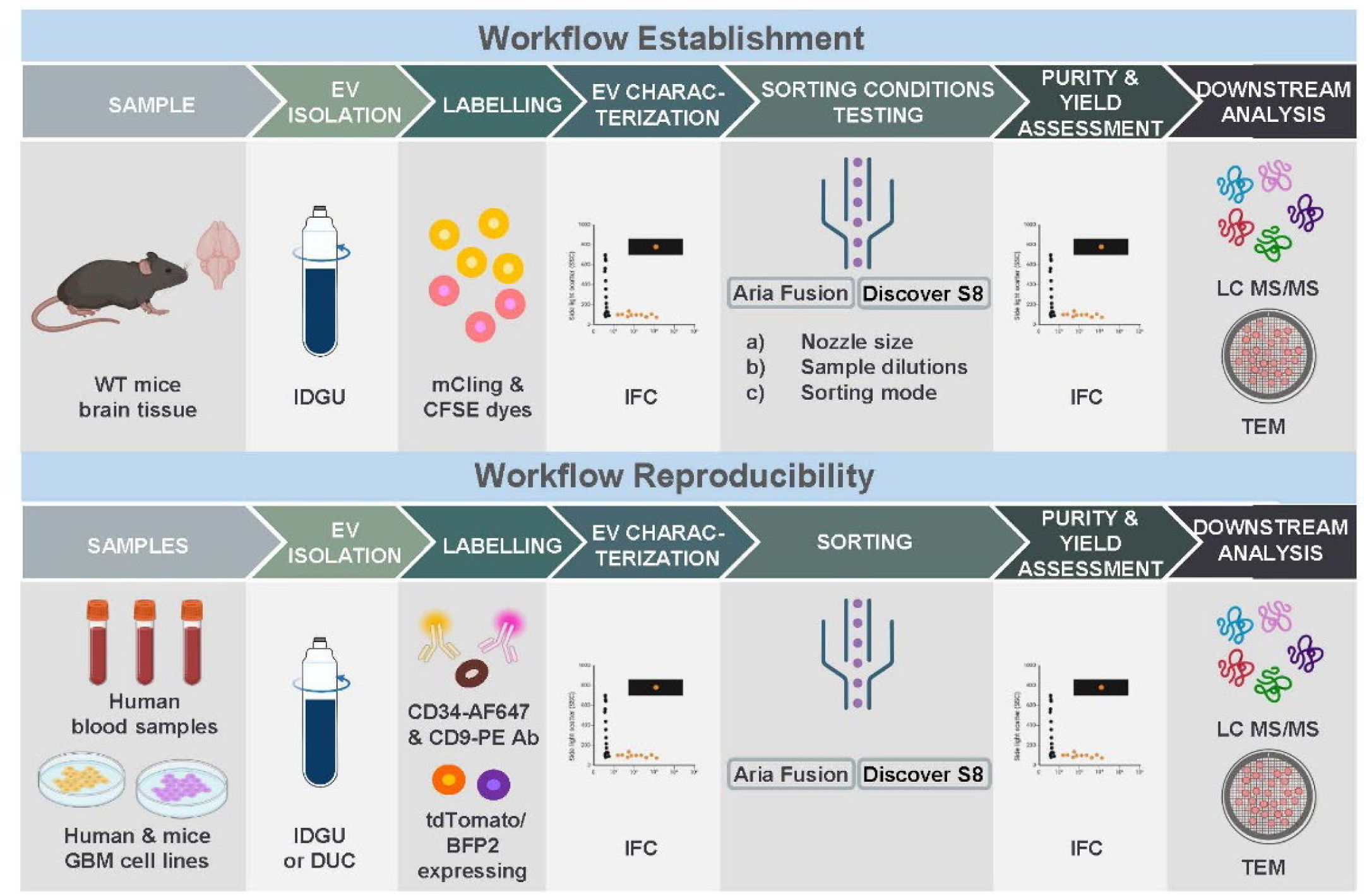
Study overview. Graphical representation of the experimental setup illustrating protocol establishment (top) and subsequent reproducibility (bottom). IDGU: iodixanol density gradient ultracentrifugation; DUC: differential ultracentrifugation; IFCM: imaging flow cytometry; LC-MS/MS: liquid chromatography/tandem mass spectrometry; TEM transmission electron microscopy.

### State-of-the-art droplet-based FACS instruments can detect particles as small as 50 nm from background noise, enabling EV sorting

A key challenge of using flow cytometric sorters for EV enrichment is the substantial size difference between cells and EVs. As a first step we assessed the resolution of the sorter to ensure sufficient sensitivity to detect particles as small as EVs. We assessed two droplet sorters with different laser power, electronics and detection systems: (I) the BD FACSAria™ Fusion (hereafter Fusion) representing a conventional state-of-the-art sorter and (II) the BD FACSDiscover™ S8 (hereafter S8), a next generation spectral sorter with imaging capabilities. We used FSC-SSC-Megamix beads and NIST beads with a range size of 100–900 nm and 50-90 nm, respectively to evaluate the lower detection limits and particle resolution capabilities of both sorting platforms for EV-sized particles. As shown in **Figure 2A** and **2B** both sorters were able to resolve bead populations of different sizes, also within the EV size range. Importantly, even the smallest bead size (50nm), was clearly distinguishable from the PBS background. However, it must be taken into account that the refractive indices of calibration beads and EVs differ substantially. Consequently bead-based measurements provide only an approximation of the sorter’s capability for the detection and resolution of EVs^34^

**Figure 2.**
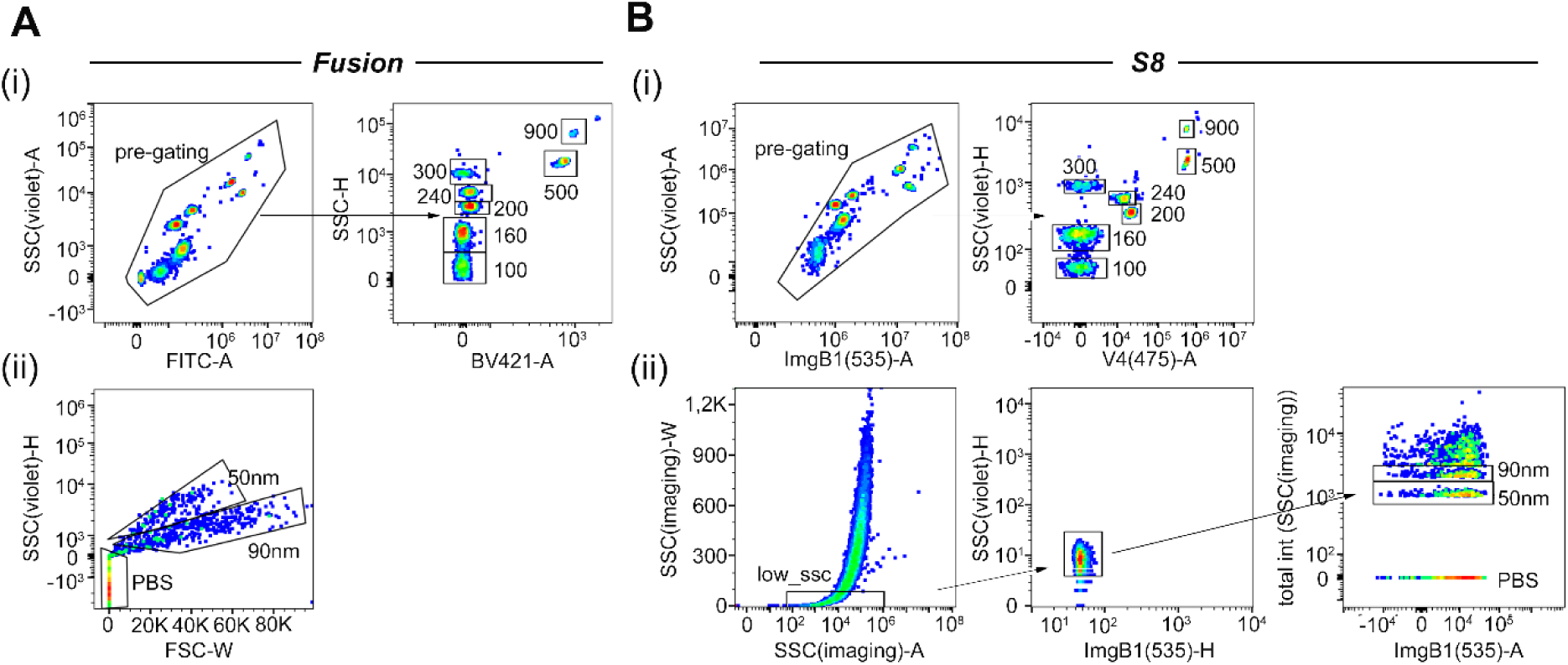
Technical adjustments enabling the use of FACS instruments for EV sorting. FSC-SSC Megamix beads (i) and NIST-beads (ii) in the size-range of EVs are successfully detected in the Fusion (A) or S8 (B).

To enable EV sorting, a tightly focused sample stream is required allowing individual EVs, rather than multiple coincident EVs (’swarming’), to be analyzed while excluding non-EV components. In addition, the transit time between the interrogation point and the droplet break-off point must be precisely controlled to ensure the accurate charging and deflection of the correct droplet into the designated collection tube. Maintaining optimal pressure conditions is also essential to minimize potential structural damage to EVs. The key adjustable parameters enabling these conditions include: (i) nozzle size, which consequently affects sheath pressure and drop drive frequency; (ii) sample dilution; (iii) sorting flow rate and core stream diameter; and (iv) sort precision mode. **Table 2** summarizes the physical principles underlying these critical adjustments for EV sorting.

As stated above, we established the sorting parameters with mouse tissue brain-derived EVs. The isolation of EVs directly from tissues has gained increasing attention in recent years^36^. Directly isolating EVs from tissue enables the study of a highly diverse EV population that retains, probably partially, the EV corona, integrates information from cell-to-cell interactions, and captures the temporal patterns of disease progression, difficult to replicate under cell culture conditions. Therefore, mouse tissue brain-derived EVs represent a highly valuable source of disease-relevant information. Mouse tissue brain-derived EVs were isolated from wild-type (WT) mouse brain through differential gradient ultracentrifugation following published protocols^22^ and recommendations^37^. For this proof-of-principle experiment, two mouse tissue brain-derived EV samples were prepared and labelled separately with fluorescent dyes before being combined, thereby generating two distinguishable EV subsets. One subset was labelled with mCling-Atto 647, a membrane-binding dye that remains associated with plasma and endocytic membranes^38^, whereas the other subset was labelled with CFSE, a cell-permeable dye that covalently binds intracellular proteins, widely used in flow cytometry. (**Figure 3A**). NTA measurements performed prior to sorting revealed that mCling⁺ and CFSE⁺ EVs exhibited modal particle sizes of 183.9 ± 23.0 nm and 146.4 ± 17.8 nm, respectively (**Supplementary Figure 2A**). Their characteristic cup-shaped morphology was confirmed by TEM **Supplementary Figure 2B)**, while expression of EV-associated tetraspanins (TSPN) CD9, CD81 and CD63 was confirmed by IFCM (**Supplementary Figure 2C-D**). The two labelled EV subsets were mixed in defined proportions of approximately 30% mCling-labelled EVs and 70% CFSE-labelled EVs (**Figure 3B**). The aim was to enrich the less abundant mCling^+^ EV population from the more abundant CFSE^+^ population.

**Figure 3.**
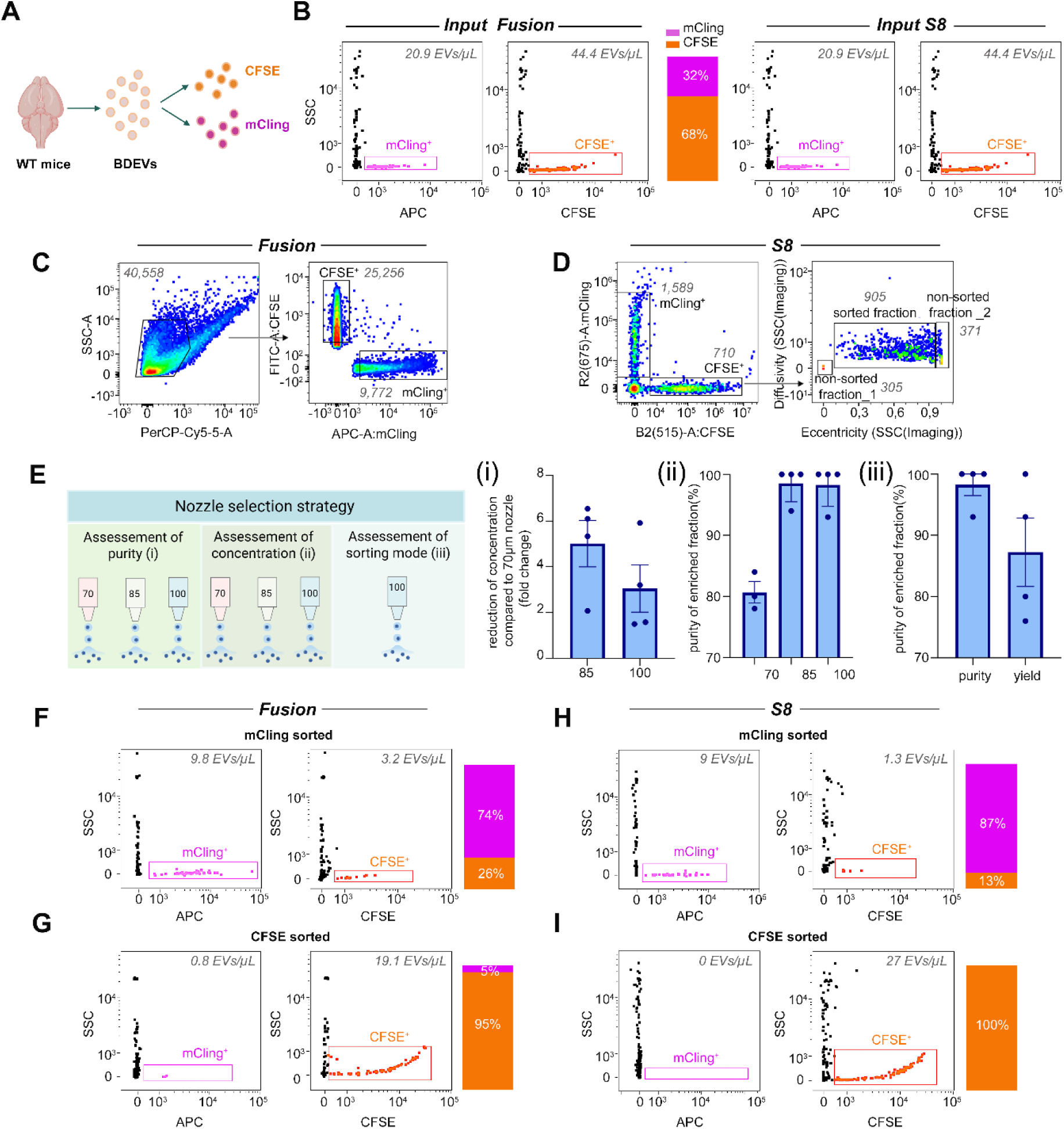
Sorting of mouse tissue brain-derived EVs. A) Experimental set up; B) IFCM analysis of the input EV samples for the sorting with the Fusion (left) or the S8 (right). Gating strategy for sorting CFSE^+^ EVs and mCling^+^ EVs using the Fusion (C) or S8 (D); E) Graphical depiction of the nozzle selection strategy. (i) Sorted EV Concentration fold change reduction using the 85µm and 100µm Nozzle compared to the 70µm Nozzle; (ii) EV enrichment by nozzle size; (iii) EV enrichment dependent on sorting mode; F) IFCM analysis of the sorted fraction when sorted for mCling^+^ EVs or CFSE^+^ EVs (G) with the Fusion; H) IFCM analysis of the sorted fraction when sorted for mCling^+^ EVs or CFSE^+^ EVs with the S8 (I).

For sorting with the Fusion, swarmed particles and non-EV-particles were excluded by first gating for low sideward scatter (SSC-A) before gating for the fluorescent signal of interest (**Figure 3C**). For sorting with the S8, several imaging features including Diffusivity, Eccentricity, Total Intensity and Radial Moment are available to support particle discrimination from background and swarmed particles^39^. We evaluated all the imaging features and selected those that allowed for the clearest visual separation of background signals and swarmed particles when examining each type of sample in the sorter. For the mouse tissue brain-derived EVs the imaging features ‘Eccentricity’ and ‘Diffusivity’ were applied. Diffusivity reflects the local concentration of signal intensity, whereas Eccentricity is defined as the ratio of the shortest to the longest axis of the detected particle (**Figure 3D; Supplementary Figure 2E**). Using this gating strategy, we next evaluated the effect of nozzle diameter on sorting performance. We evaluated three nozzle sizes: 70 µm (available only in the Fusion sorter), 85 µm, and 100 µm. The highest concentration was obtained with the 70 µm nozzle, but the highest purity was achieved with the 85 µm and 100 µm nozzles (**Figure 3E**). For subsequent experiments we selected the 100 µm nozzle for two reasons: (i) it provided a better enrichment in terms of purity than the 70 µm nozzle while not showing substantial differences compared to the 85µm nozzle in this regard and (ii) it operates at lower pressure than the 85µm nozzle, thereby reducing shear stress and minimizing the risk of EV damage during sorting (**Table 2**). Sample dilution conditions were also optimized to ensure clear resolution of the particle populations while minimizing swarm events. Appropriate sample dilution should be verified prior to sorting as insufficient dilution leads to swarm detection artefacts (**Supplementary Figure 3,** shown for S8). Regarding flow rate, we used the minimum setting (1) for both sorters to maintain a narrow core stream, thereby maximizing the probability that singular EVs pass through the laser interrogation point (**Table 2**). Finally, we assessed whether the choice of sorting mode (purity vs. yield) influenced enrichment in terms of purity (**Figure 3E**). No statistically significant differences in enrichment were observed between these two sorting modes; however the variability was substantially higher when using the sorting mode yield. Therefore, we proceeded with purity as the default sorting mode for subsequent experiments. The selected features for EV sorting are summarized in **Table 2**.

### Low abundant subsets of fluorescently labelled tissue-derived EVs can be enriched with a purity between 70% to almost 90% depending on the sorter

For fluorescently labelled mouse tissue brain-derived EVs, the selected technical sorting parameters and gating strategy resulted in an enrichment of the less abundant mCling^+^ EVs population with a purity up to 74% and of the more abundant CFSE^+^ EVs population up to 96% as determined via IFCM following sorting with the Fusion (**Figure 3F, G**). Using the S8, the enrichment led to a purity of mCling^+^ EVs population up to 87% and CFSE^+^ EVs population up to 100% (**Figure 3H, I**). Controls according to MIFlowCyt-EV guidelines^40^ are summarized in **Supplementary Table 2** and shown in **Supplementary Figures 4 and 5.** Additionally, serial dilutions are shown in **Supplementary Figure 6.** To ensure the reproducibility of our results, we provide information on the position of the reference beads relative to the applied gates(**Supplementary Figure 7)**. This allows future researchers to position their own gates relative to the same beads, enabling successful EV sorting.

Here, we show that EVs from demanding isolation protocols labelled with intraluminal and intramembrane dyes, can be successfully sorted, providing a valuable tool for investigating EV biology in this context.

### Rare EV subsets in blood can be enriched with a purity of nearly 90% and 100% depending on the sorter

Next, we selected blood-circulating EVs to demonstrate that our EV enrichment workflow is also applicable to this sample type, which is of particular interest as serum represents the most commonly investigated source of biofluid-derived EVs^41^. However, serum remains a challenging source for EV analysis because of the high abundance of lipoproteins and platelet-derived particles. EVs were isolated from serum samples using differential ultracentrifugation. EV isolation was validated using NTA, TEM and IFCM as before. The modal diameter of the EVs to be sorted was 124.2 nm (+/- 11.4 nm) and TEM images showed the characteristic cup-shape EV morphology **(Supplementary Figure 8A and 8B).** The presence of established TSPNs as well as the absence of serum-derived contaminants was evaluated with IFCM (**Supplementary Figure 8C-E).** EVs were then labelled with fluorophore-conjugated antibodies against CD34(-AF647), an endothelial marker and CD9(-PE), a well-established marker for EVs (**Figure 4A**). Since EV populations of clinical interest are typically present at low frequencies within a large background of unrelated circulating EVs, we established a defined rare-population model. EVs isolated from one blood sample were labelled with CD9 and served as the background population, whereas EVs isolated from a second blood sample were labelled with CD34 and used as the target population. Low amounts of CD34+ EVs were then spiked into the larger CD9+ EV pool, resulting in input samples containing 7% CD34+ EVs for sorting with the Fusion and 12% CD34+ EVs for sorting with the S8 (**Figure 4B**). This approach generated a controlled and predefined rare EV population, thereby mimicking the challenge of isolating low-abundance EV subsets, such as tumor-derived EVs, from complex biological samples.

**Figure 4.**
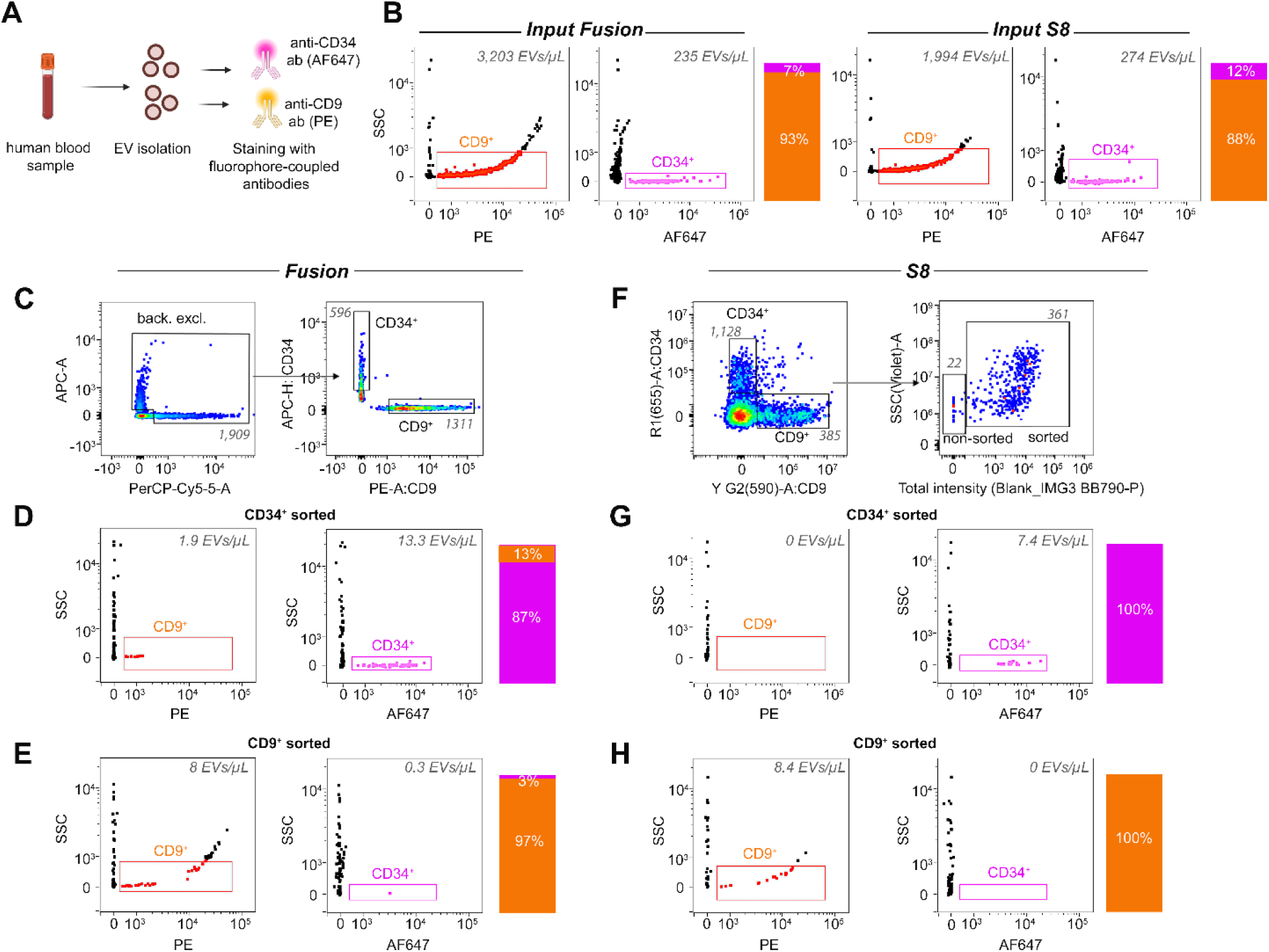
Sorting of human blood-circulating EVs. A) Experimental set up; B) IFCM analysis of the input for sorting with the Fusion (left) and the S8 (right); C) Gating strategy for sorting CD34^+^ EVs and CD9^+^ EVs on the Fusion. IFCM analysis of the sorted fraction with the Fusion for CD34^+^ EVs (D) or CD9^+^ EVs (E); F) Gating strategy of sorting for CD34^+^ EVs and CD9^+^ EVs on the S8. IFCM analysis of the sorted fraction with the S8 for CD34^+^ EVs (G) CD9^+^ EVs (H).

Following evaluation of serial dilutions (**Supplementary Figure 6B**), highly diluted samples were used on both sorters, yielding target population event rates of 6–9 events/sec (361–596 events/min). This allowed clear resolution of non-swarm events within the sorting gates (Figure 4C, F). For sorting with the Fusion, the first gate applied was intended to distinguish fluorescently labelled EVs from background, followed by subsequent gating for the fluorescent populations of interest (**Figure 4C**). We obtained an enrichment of nearly 90% purity for CD34^+^ EVs and nearly 100% purity when sorted for CD9^+^ EVs as validated by IFCM (**Figure 4D, E**). For sorting with the S8, the “Total Intensity” feature - which integrates all intensity values of all pixels for one parameter – was gated against the SSC-A (**Figure 4F**). This approach enabled the identification of events exhibiting high fluorescence intensity values and their discrimination from background contaminants (**Supplementary Figure 8F)**. Additional imaging features, including ‘Eccentricity’, were also evaluated; however, the best separation of the target population was achieved using the ‘Total Intensity’ feature. Using this gating strategy, we achieved an enrichment of both sorted fractions reaching up to 100% (**Figure 4G, H**). Controls according to MIFlowCyt-EV guidelines^40^ are summarized in **Supplementary Table 2** and the position of the reference beads relative to the gates to ensure reproducibility are provided in **Supplementary Figures 9-11**.

Our results show that despite the challenges associated with blood-circulating EVs, high enrichment rates up to 100% purity are achieved when labelling EVs with antibody-coupled fluorophores and even if the population of interest represents around 10% of the overall EV population. The S8 imaging features employed here allowed clear distinction from background signals and swarmed particles. This opens a unique opportunity to analyze rare EV populations in blood for diagnostic and therapeutic purposes.

### Enrichment of cell culture-derived EV subpopulations resulted in more than 90% purity in both sorters

Lastly, we validated our sorting approach using EVs from glioma cell lines, which are widely used models for studying brain tumor biology ^42^.

The handling of cell culture-derived EVs is generally considered as straightforward since conditioned media is a relatively uncontaminated EV source, compared to biofluids such as blood^43^.

EVs were isolated by differential ultracentrifugation of conditioned media from the human glioma-derived cell line (NCH1681 line) genetically modified to express palm-tdTomato, and from the murine glioma cell line (Mut3) transduced to express BFP2 (Blue Fluorescent Protein 2, from now onwards BFP2) (**Figure 5A**). EVs were characterized by NTA, TEM and IFCM. tdTomato^+^ EVs had a modal size of 153.4nm (+/- 6.6 nm) and BFP2^+^ EVs had a modal size of 175.7nm (+/- 5.2 nm) (**Supplementary Figure 12A**). TEM showed the expected cup-shape morphology (**Supplementary Figure 12B**) and IFCM showed the presence of TSPNs confirming successful EV isolation (**Supplementary Figure 12C-D**). For the sort we prepared an input mixture consisting of approximately 80% tdTomato^+^ EVs and 20% BFP2^+^ EVs for sorting with the Fusion and approximately 65% tdTomato^+^ EVs and 35% BFP2^+^ EVs for sorting with the S8, as determined by IFCM (**Figure 5B**).

**Figure 5.**
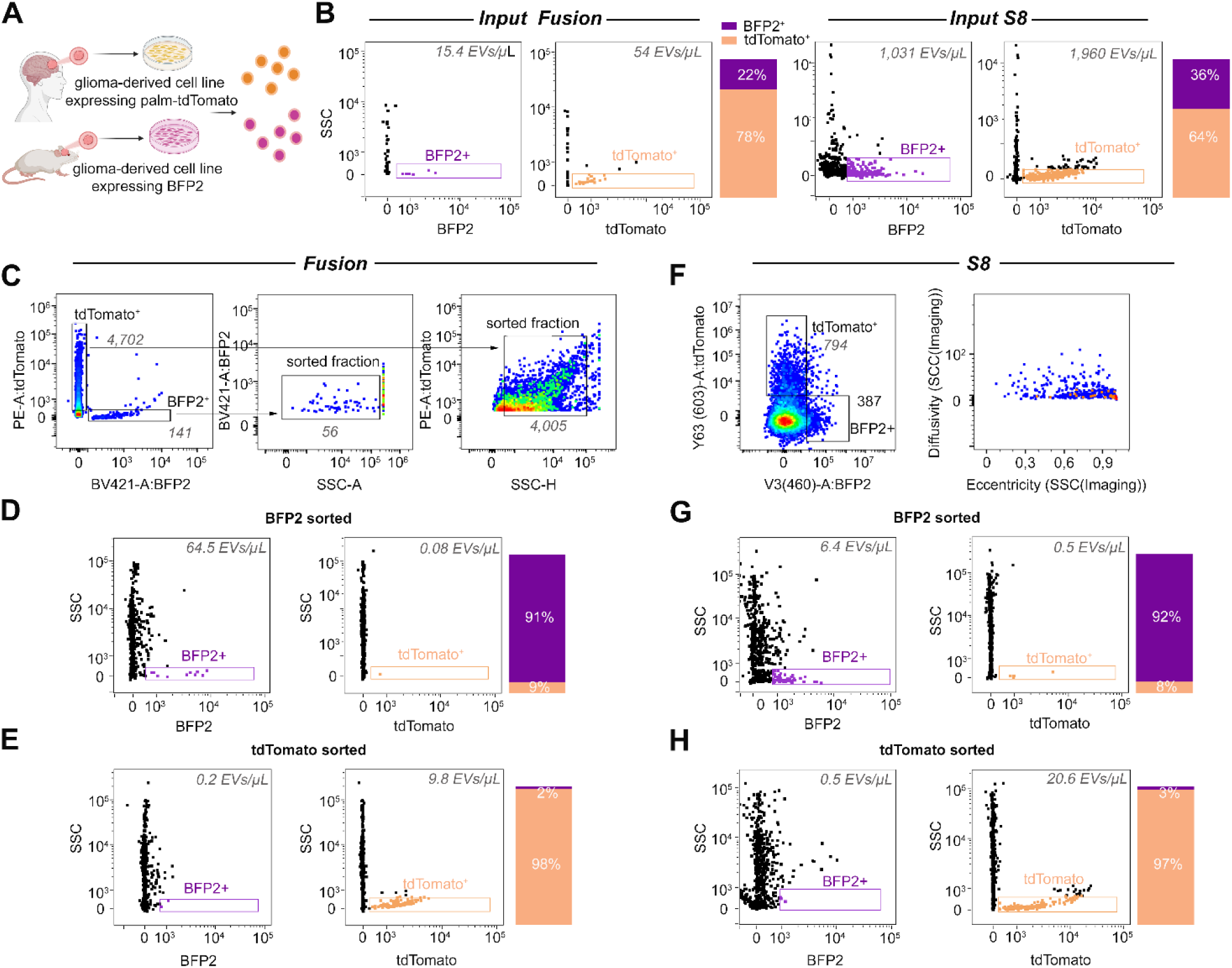
Sorting of glioma cell culture-derived EVs. A) Experimental set up; B) IFCM analysis of the input for the sorting with the Fusion (left) and the S8 (right); C) Gating strategy for sorting tdTomato^+^ EVs and BFP2^+^ EVs on the Fusion. IFCM analysis of the sorted fraction with the Fusion for BFP2^+^ EVs (D) or tdTomato^+^ EVs (E).IFCM analysis of the sorted fraction with the S8 BFP2^+^ EVs (G) or tdTomato^+^ EVs (H).

As cultured media from glioma stem-cells was highly enriched in EVs **(Supplementary Figure 12A),** following evaluation of serial dilutions **(Supplementary Figure 6C**) high dilutions were selected, resulting in event rates of 65-85 events/sec (4,000-5,000 events/min) within the sorting gates for both sorters (**Figures 5C, F**). As before, for sorting with the Fusion, swarmed EVs and unspecific signals were excluded by gating exclusively for events with SSC-A (**Figure 5C**). This resulted in an enrichment of BFP2^+^ EVs with a purity up to 91% and of tdTomato^+^ EVs up to 98% as confirmed by IFCM (**Figure 5D,E**). For sorting with the S8 and in contrast to the other preparations, none of the evaluated image features provided additional separation of cell cultured-derived EVs from background signals, as shown for the parameters Diffusivity and Eccentricity in **Figure 5F**. This may reflect the relatively low level of contaminating particles present in conditioned medium-derived EV preparations, limiting the additional discriminatory power provided by the imaging features. Consequently, positive fluorescence signals without further gating were selected for sorting. This resulted in post-sort purities approaching 100% for both BFP2^+^ and tdTomato^+^ EVs (**Figure 5G, H**). Corresponding controls as suggested by the MIFlowCyt-EV guidelines^40^ can be found in **Supplementary Figure 13** and the position of the reference beads relative to the gates are shown in **Supplementary Figure 14**.

In summary, endogenous fluorescent reporter proteins such as tdTomato and BFP2 generated emissions that were readily distinguishable from background. Both sorting platforms achieved high-purity enrichment of the target EV population, exceeding 90% purity while maintaining minimal contamination. Notably, this level of enrichment was achieved irrespective of the abundance of the target population in the input sample.

To demonstrate the robust reproducibility of our sorting strategy, we performed several independent sorting experiments on different days using samples prepared in separate batches. In all cases, we obtained highly consistent results (**Supplementary Figure 15**).

### Post-sorted EVs maintain their integrity and stability enabling downstream analysis

After sorting, we evaluated whether the isolated EVs retained key properties including integrity and stability. Specifically, we performed NTA, TEM and LC–MS/MS tandem mass spectrometry. NTA was used to measure recovery and size distribution after sorting. Recovery rates varied substantially across sample types. For mouse tissue brain-derived EVs, recovery of the target population was high, with only minimal deviation between input and recovered EV numbers (**Supplementary Figure 16A**). In contrast, recovery of blood-circulating EVs and cell culture-derived EVs was substantially lower, with approximately one in fifteen target EVs being recovered (**Supplementary Figure 16 B, C**). Given the limited sorting duration (1-2h) applied in this study, it is possible that longer sorting times could further improve recovery. NTA analysis further indicated the absence of size-dependent selection bias across all the EV sample types (**Supplementary Figure 17**). It should be noted, however, that post-sort EV concentrations were relatively low, likely reducing the sensitivity and accuracy of the size measurements.

TEM analysis of sorted cell culture-derived EVs confirmed the presence of intact EVs following sorting (**Figure 6A**). Interestingly, we observed that successful TEM visualization required the use of highly positively charged grids as EVs were not detectable when using negatively charged grids. We hypothesize that the electrical charge applied during sorting altered EV-grid interaction reducing their binding capacity on negatively charged surfaces.

**Figure 6.**
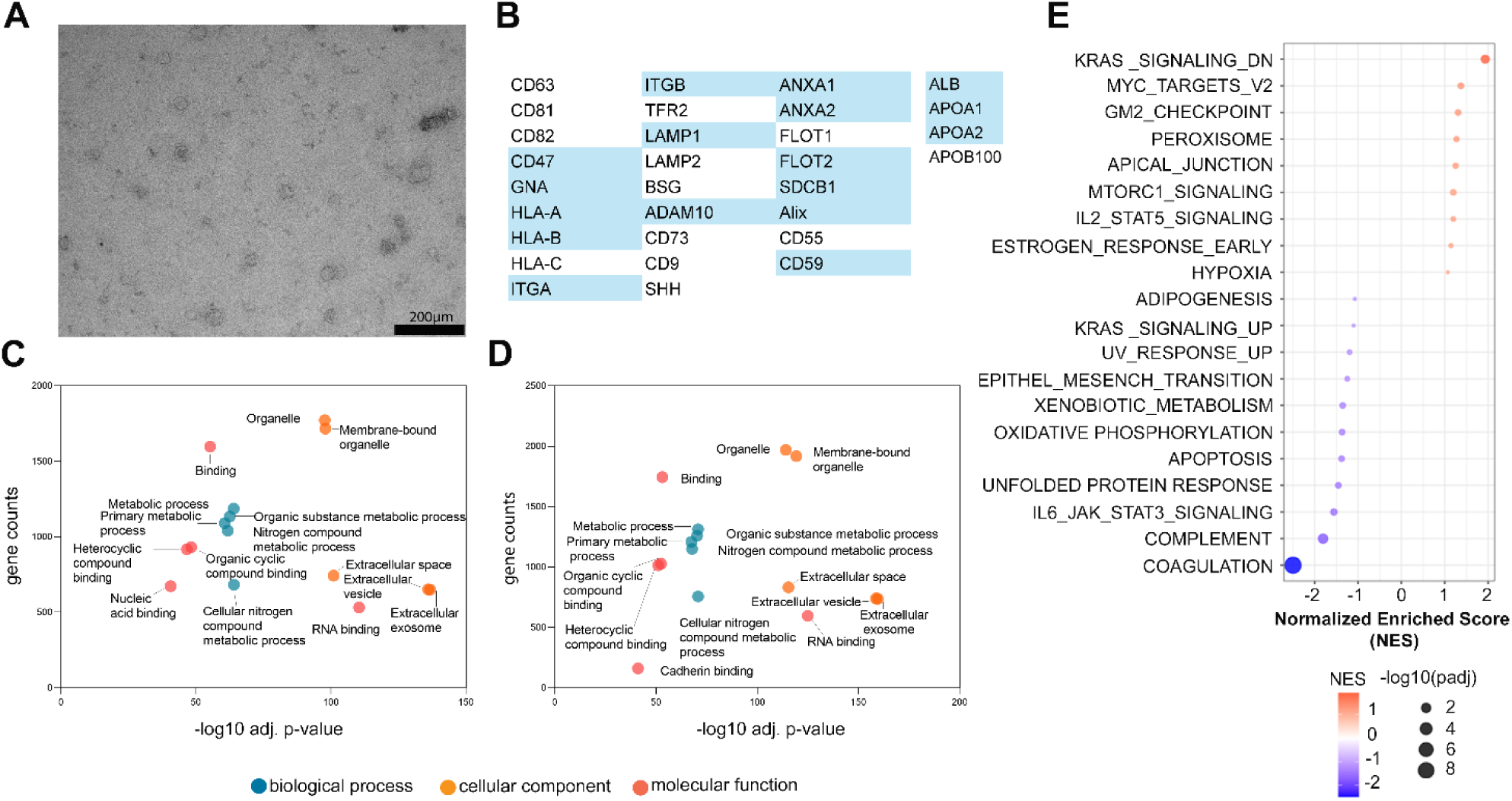
Effective EV population recovery after sorting. A) TEM of BFP2^+^ EVs after sorting; B) Common EV markers as well as contaminants identified among the proteins after sort (blue = detected, white = not detected); C) Gene ontology analysis of CD34^+^ EVs after sorting with the Fusion; D) Gene ontology analysis of CD34^+^ EVs after sort with S8; E) Gene Set-Enrichment-Analysis of CD34^+^ EVs before and after sort.

Lastly, to assess the suitability of sorted EVs for downstream applications, sorted human blood-circulating EVs and mouse tissue brain-derived EV samples were analyzed using LC-MS/MS. Proteomic analysis of all sorted CD34+ EVs, -sorted from EVs initially isolated from 250µl serum- shows that this input volume is sufficient to detect approximately 2,000 proteins (**Figure 6B, Supplementary Table 3**). Among the identified proteins were established EV-associated markers, including Flotilin, Annexins, and Alix (**Figure 6B**). Proteins such as albumin or apolipoproteins were also detected. While these proteins have been traditionally considered contaminants in EV preparations, accumulating evidence suggest that they may play functional roles as components of the EV corona^44^. Therefore, their persistence after sorting may not necessarily reflect incomplete purification but could instead be related to their association with EVs^45,46^. Gene ontology analysis conducted after the sort (**Figure 6D, E**), showed presence of EV-related terms across three GO categories, such as ‘Extracellular exosome’, RNA-binding,’ and ‘Metabolic process’. In addition, gene set enrichment analysis (GSEA) comparing pre- and post-sort human blood EV samples revealed an increased enrichment for cell type–specific pathways, whereas coagulation- and complement-associated protein signatures were diminished (**Figure 6F)**. Likewise, proteomics of CFSE+ EVs confirmed successful isolation, identifying EV-specific markers (**Supplementary Figure 18**).

## DISCUSSION

To our knowledge, we provide for the first time a workflow for selective EV sorting using both conventional and spectral FACS instruments across diverse EV species, source, size, isolation methods, and fluorophore-labelling strategies. This workflow enables high-purity enrichment of EVs, including low-abundance populations (<10%), while preserving stability and integrity for downstream applications. Among the tested instrument parameters, the nozzle diameter had the strongest influence on sorting performance. Although the 70 µm nozzle resulted in higher sorting concentrations, its enrichment performance in terms of purity was lower than that achieved with the 85 µm and 100 µm nozzles. This observation may reflect differences in droplet formation dynamics, which are known to affect sorting accuracy by influencing whether detected events are correctly directed to the intended collection stream. When sorting using the 100 µm nozzle (selected based on low operating pressure), enrichment across all sample types reached up to 100%. Larger nozzle sizes may also reduce the risk of EV structural damage, thereby improving the practical suitability of the method for maintaining EV integrity. Consistent with this interpretation, Higginbotham et al. reported the integrity of sorted EVs by dSTORM microscopy when sorting with a 100µm nozzle^9^. The choice of nozzle size involves a trade-off between droplet size, which affects the likelihood of single-EV occupancy, and avoiding excessive pressure, which could compromise EV integrity. With the 100 µm nozzle, this balance was achieved. It must be noted that larger nozzle sizes also increase the droplet volume, which can lead to larger collection volumes and lower final sample concentration. Depending on the intended downstream application, an additional concentration step after sorting may therefore be required.

A second critical parameter was sample dilution. Sufficient dilution is required to enable reliable single-particle measurements and to minimize coincidence and “swarming” of multiple events. Inadequate dilution might compromise separation quality. Previous work from Kormelink et al. using serial dilutions of 100 nm reference beads, liposomes, and EVs showed that high dilution is essential not only to prevent coincidence events but also to reduce excessive light scattering and fluorescent intensities^35^. Together, these results highlight the importance of optimizing nozzle size and sample dilution as, different to cell sorting, even small deviations from single-particle occupancy might increase the probability of multiple vesicles sorting, compromising purity.

Regarding the sorting mode, prior studies have shown that purity sorting mode often results in higher enrichment rates and lower variability than yield mode^47^. We observed similar trends with yield mode resulting in a higher variability, leading us to conclude that, as for cell sorting, purity mode is the most adequate mode for EV sorting.

Previous studies have reduced background signals by introducing additional EV washing steps, such as size-exclusion chromatography (SEC) after staining, to remove excess fluorophores and contaminants. In the present workflow, effective discrimination of background events was achieved through optimized gating strategies. This avoids additional processing steps that may result in EV loss while still enabling high-purity enrichment. For the Fusion, high-purity sorting relied on appropriate gating using scatter and fluorescence information (e.g., low SSC-A followed by fluorescence-based discrimination for EV-labelled targets). This two-step approach effectively reduced false positives, which is consistent with the fact that EV-sized background events can show overlapping scatter/fluorescence patterns that are not distinguishable without additional context. For the S8, we leveraged on image-based spectral imaging features to exclude background signals and contaminants. Different S8 imaging features performed differently depending on the sample type and labelling strategy. For example, for mouse tissue brain-derived EVs, optimal visual separation from background was achieved with Eccentricity and Diffusivity, whereas for blood-derived EV the Total Intensity feature provided the best separation. These differences likely reflect variations in contaminant composition across EV sources. Serum preparations contain abundant lipoproteins and platelet-derived particles with distinct imaging signatures compared to more controlled sources such as conditioned media, where no imaging features were needed for separation from background particles. In the future, advances in high-sensitivity detectors, optics and in image-based gating strategies, potentially combined with machine learning approaches may further improve the resolution and discrimination of even rarer EV subsets.

There are limitations to our workflow. The highest enrichment was achieved using the 100 µm nozzle, however this nozzle size led to a lower overall concentration compared with, for example, the 70 µm nozzle. Also, recovery rates varied across sample types: mouse tissue brain-derived EVs showed the best performance, with minimal losses, whereas greater losses were observed for blood-circulating EVs and cell culture–derived EVs. Extending the sorting time beyond the one hour used here would likely increase the recovery. Importantly, despite the reduced recovery relative to the input fraction, the enriched material remained sufficient for downstream applications. In particular, TEM confirmed intact EV morphology, and LC-MS/MS proteomics identified thousands of proteins with EV-enriched profiles. The detection of established EV markers in the enriched fractions, together with EV-associated Gene Ontology (GO) terms, supports the conclusion that this sorting workflow preferentially enriches bona fide EVs rather than fluorescent artefacts or vesicular debris.

An additional consideration is that both instruments evaluated in this study are droplet-based sorters. The observed enrichment performance may therefore depend, at least in part, on the physical principles of droplet generation and electrostatic deflection used in these systems. Whether comparable enrichment efficiencies can be achieved using alternative sorting technologies, such as microfluidic or acoustic-based sorters, remains to be determined. Future studies should therefore evaluate the applicability of this workflow across non-droplet sorting platforms.

The primary goal of this study was to demonstrate the applicability of the workflow across different sample types and on to two droplet-based FACS platforms with different detection paradigms. We did not aim to perform a standardized cross-platform comparison of the instruments and therefore did not include arbitrary unit normalization. Instead, we aimed to show that the workflow enables reliable, high-purity EV enrichment on both platforms. Future studies aimed at inter-laboratory comparisons and cross-platform standardization should incorporate normalization to arbitrary units to facilitate direct comparison of results^48^.

Collectively, our workflow enables FACS-based EV sorting as a practical and reproducible tool for investigating EV heterogeneity across diverse sources and labelling strategies. By enabling the enrichment of defined EV populations while preserving compatibility with downstream analyses, this approach provides a valuable tool for both fundamental EV biology and translational research applications.

### Geolocation information

Human blood and mouse brain tissue samples were collected in Hamburg, Germany. Cell culture experiments and EV analyses were performed at the University Medical Center Hamburg-Eppendorf, Hamburg, Germany.

## Supporting information

MiBloodEV supplementary data

MIFlowCytEV supplementary data

## Acknowledgments

We are grateful to Dr. Christel Herold-Mende (Heidelberg University Hospital, Heidelberg, Germany) for providing NCH1681 cell line, as well as Dr. Sean Lawler (Brown University, USA) for the mut3 mouse glioma line. We are also grateful to Dr. Xandra Breakefield lab (MGH, Harvard, Boston, USA) for providing us with the palm-tdTomato and palm-mTagBFP2 plasmids. We thank the Core Facility Mass Spectrometric Proteomics as part of the Technology Platform Mass Spectrometry (TPMS) at University of Hamburg (UHH) and University Medical Center Hamburg-Eppendorf (UKE) for support with mass spectrometric measurements and analysis funded by the Deutsche Forschungsgemeinschaft (DFG, German Research Foundation) – 518551069. The flow cytometer and sorter used in this study were provided by the Cytometry and Cell Sorting Core Facility, Deańs Office for Research, University Medical Center Hamburg-Eppendorf. We further acknowledge the support of the staff of the Neurosurgery Department, the Neurology Department (ERSI lab), and the Laboratory of Experimental Feto-Maternal Medicine.

Illustrations were created with BioRender (https://BioRender.com).

## Author contributions

Conceptualization: I.G., B.P., J.B., A.R, A.S-S.; Methodology: I.G., J.B., A.R, K.L.; Investigation: I.G., B.P, J.B., A.R, A.S-S., S.B., H.U., B.S., C.U., C.L.M., K.R.; Visualization and presentation of data: I.G., B.P.; Project funding acquisition: A.D., P.A., F.L.R., T.M. B.P.; Project administration: I.G.; Supervision: B.P.; Writing—original draft: I.G., B.P.; Writing—review & editing: all authors.

## Data availability statement

all the data is available upon request. The mass spectrometry proteomics data have been deposited to the ProteomeXchange Consortium via the PRIDE partner repository with the dataset identifier PXD069156.

## Ethics approval statement

For <u>human samples (blood)</u>: All study subjects signed informed consent forms. The PRINCE study protocol was approved by the e<ythics committee of the Hamburg Chamber of Physicians under the license number PV 3694 and was conducted according to the Declaration of Helsinki for Medical Research involving Human Subjects. For <u>mouse samples (brain)</u>: Animal experiments were approved by the local animal care committee (Behörde für Justiz und Verbraucherschutz of the Freie und Hansestadt Hamburg, project number N105/2023 and ORG1055) and in compliance with the guidelines of the animal facility of the University Medical Center Hamburg-Eppendorf.

