## Supplementary material for "Enrichment of Extracellular Vesicles Subsets from Diverse Biological Sources Using Conventional and Image-Based Fluorescence Activated Sorting": Final_Supplementary Data Graf et al 2026.pdf

### SUPPLEMENTARY FIGURE 1

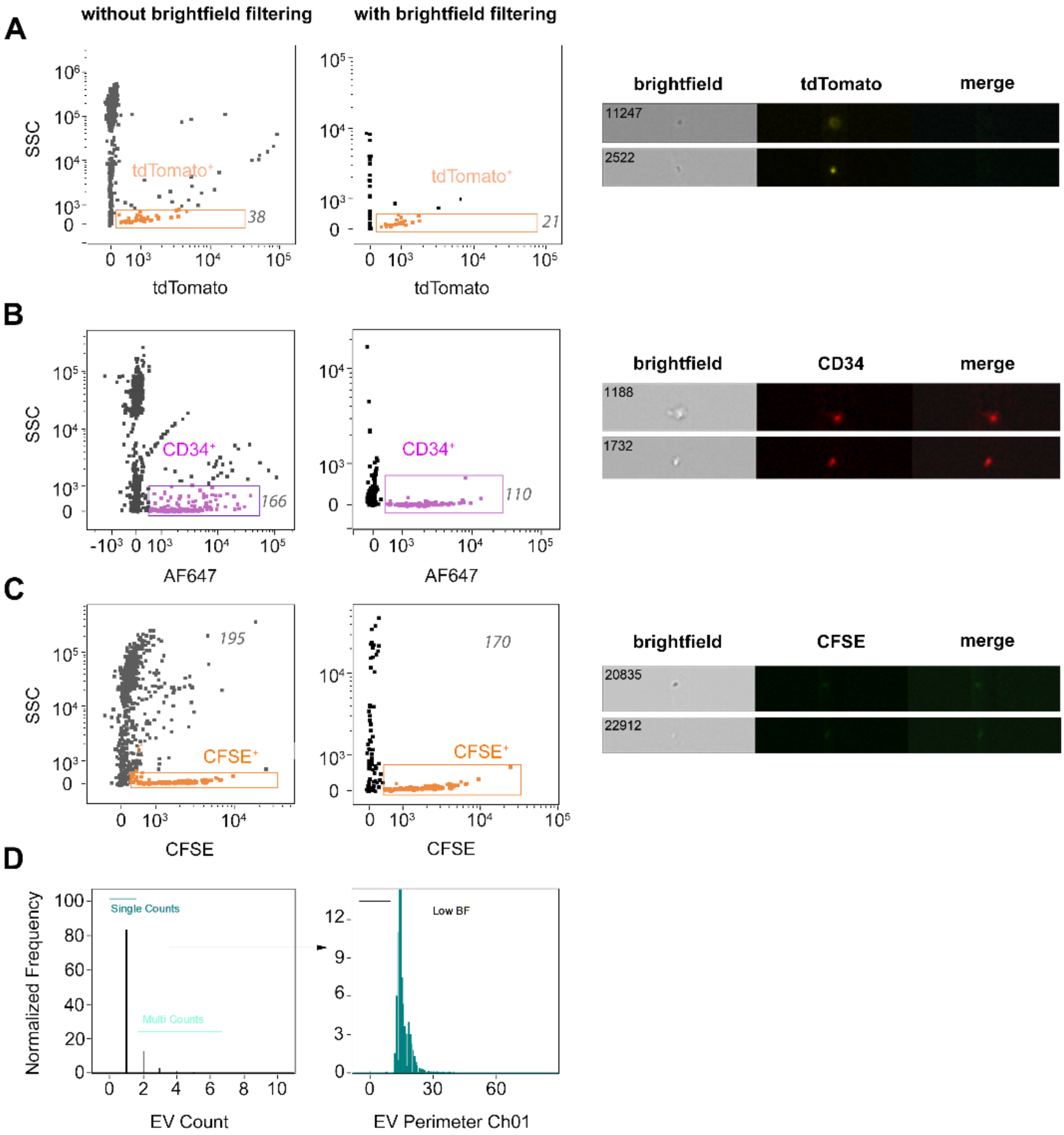


**Supplementary Figure 1. IFCM allows highly sensitive detection of EVs**. A) Gating for tdTomato⁺ EVs without (left) and with (right) imaging brightfield filter. Representative images of events included in the EV gate when no imaging filter is applied are shown; B) Gating for

CD34⁺ EVs (AF647 channel) without (left) and with (right) imaging brightfield filter. Representative images of events included in the EV gate when no imaging filter is applied are shown; C) Gating for CFSE⁺ EVs without (left) and with (right) imaging brightfield filter. Representative images of events included in the EV gate when no imaging filter is applied are shown; D) gating strategy applied to all IFCM analyses to exclude doublets and swarmed particles using the count filter, and to remove false-positive particles using the brightfield imaging filter.

**SUPPLEMENTARY FIGURE 2**

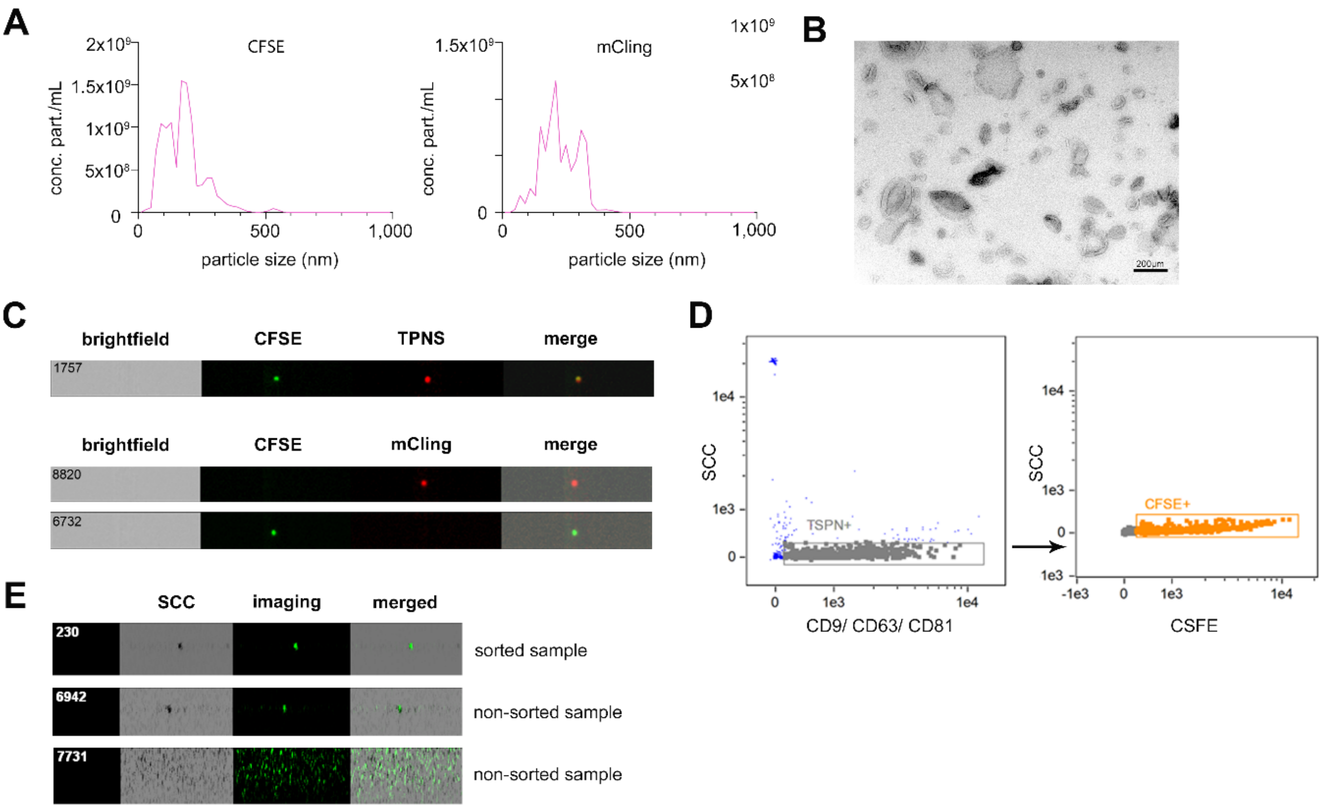


**Supplementary Figure 2**. **Characterization of mouse brain tissue-derived EVs**. A) NTA of CFSE⁺ EVs (left) and mCling⁺ EVs (right) before sorting; B) TEM of EVs before sorting; C) IFCM characterization of CFSE⁺ and mCling⁺ EVs before sort including tetraspanins (TSPN) as EV markers (top) and without TSPN (bottom); D) IFCM plots showing the co-expression of TSPN (left, CD9/CD63/CD81) and CFSE⁺ (right) on EVs before sort; E) IFCM analysis of representative images of detected events within the 'sort' and 'non sort' gates of the S8 sorter corresponding to Figure 3D.

**SUPPLEMENTARY FIGURE 3**

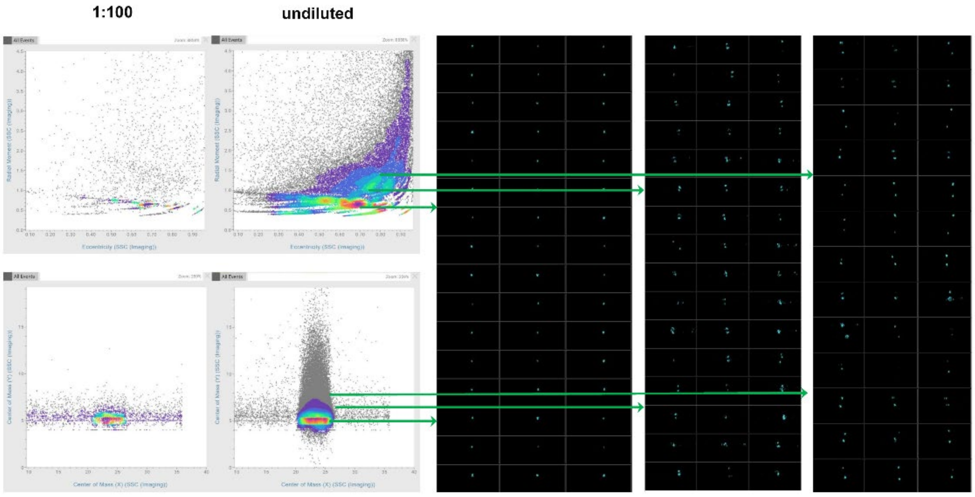


**Supplementary Figure 3. Swarming artefacts depending on dilutions shown for S8.** Representative user interface plots acquired during sorting, based on different imaging features. Left, CFSE⁺ and mCling⁺ samples at a 1:100 dilution or undiluted sample. On the right, corresponding images illustrating swarm formation in the less diluted sample, characterized by the simultaneous detection of multiple particles. This phenomenon is electronically visible by increased radial moment values of the SSC (Imaging) parameter).

### SUPPLEMENTARY FIGURE 4

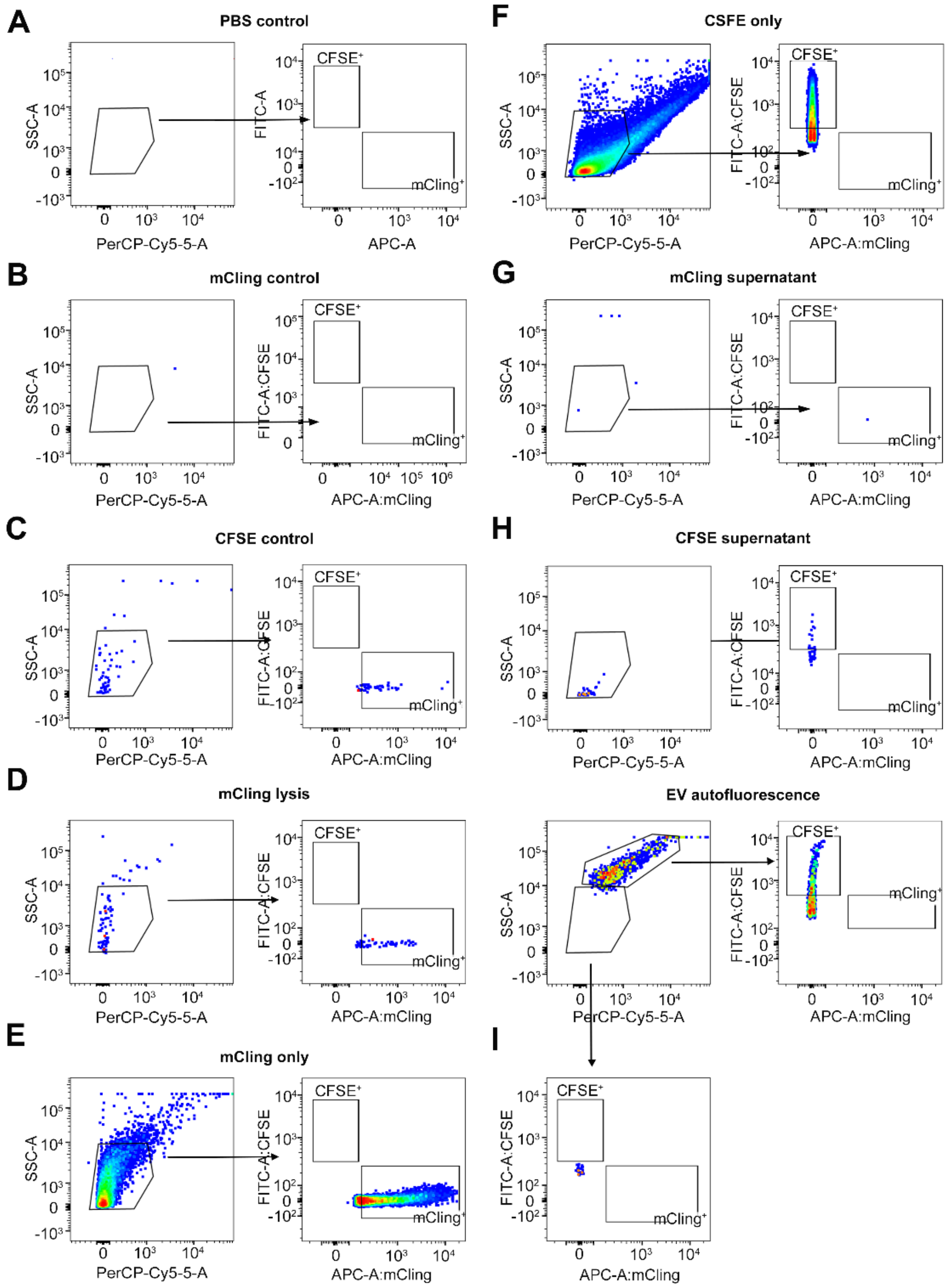


**Supplementary Figure 4. MIFlowCyt-EV sorting controls of brain tissue-derived EVs analyzed using the Fusion sorting gates**. All MIFlowCyt-EV control samples were analyzed using the same gating strategy that was applied for EV sorting. A) PBS control; B) mCling$^+$ EVs procedural control; C) CFSE$^+$ EVs procedural control; D) mCling$^+$ EVs lysis control; E) mCling$^+$ EVs single staining control; F) CFSE$^+$ EVs single staining control; G) mCling$^+$ EVs buffer with reagent control (supernatant control); H) CFSE$^+$ EVs buffer with reagent control (supernatant control); I) EVs only (autofluorescence) control.

### SUPPLEMENTARY FIGURE 5

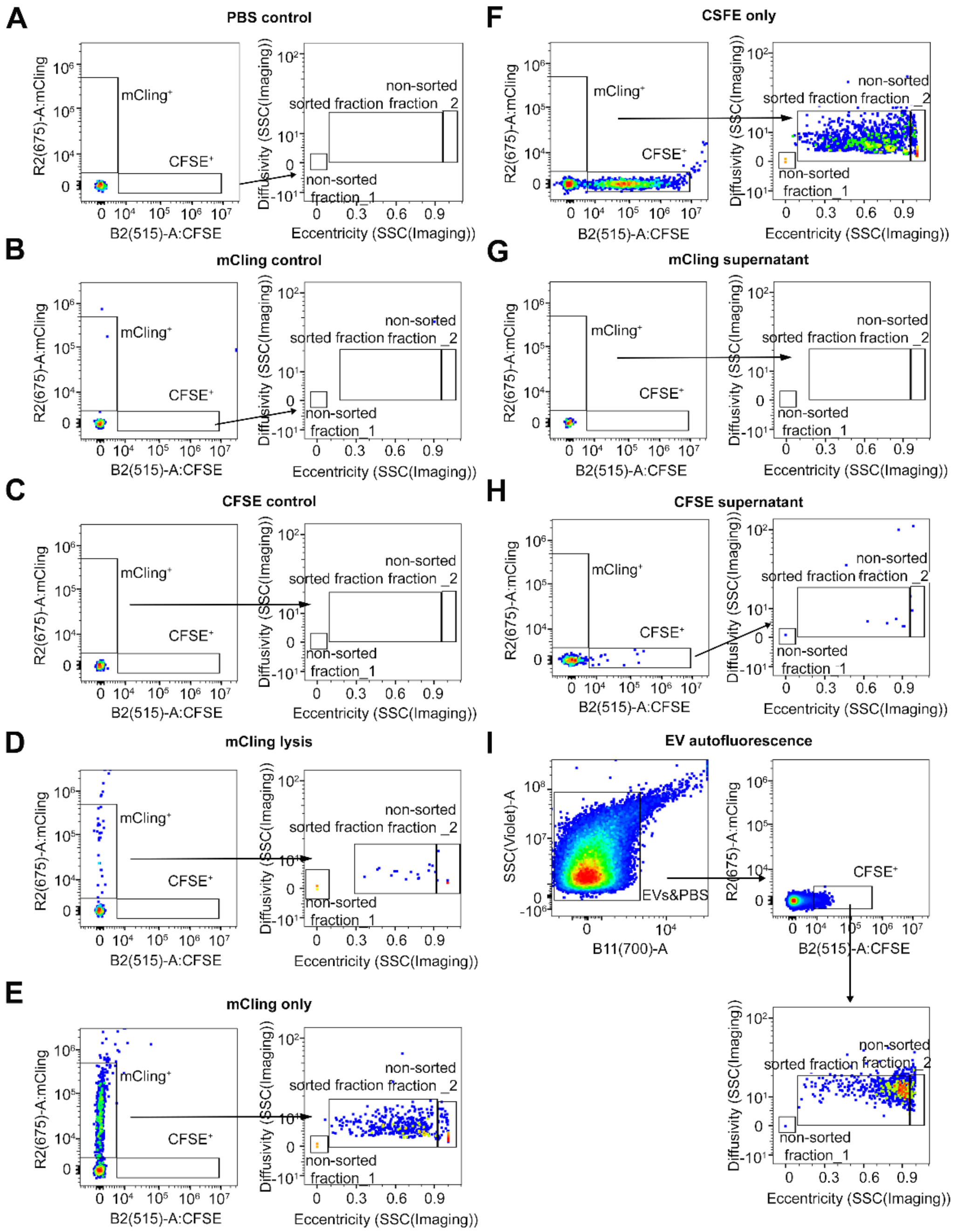


**Supplementary Figure 5. MIFlowCyt-EV sorting controls of brain tissue-derived EVs analyzed using the S8 sorting gates**. All MIFlowCyt-EV control samples were analyzed using the same gating strategy that was applied for EV sorting. A) PBS control; B) mCling$^+$ EVs procedural control; C) CFSE$^+$ EVs procedural control; D) mCling$^+$ EVs lysis control; E) mCling$^+$ EVs single staining control; F) CFSE$^+$ EVs single staining control; G) mCling$^+$ EVs buffer with reagent control (supernatant control); H) CFSE$^+$ EVs buffer with reagent control (supernatant control); I) EVs only (autofluorescence) control.

### SUPPLEMENTARY FIGURE 6

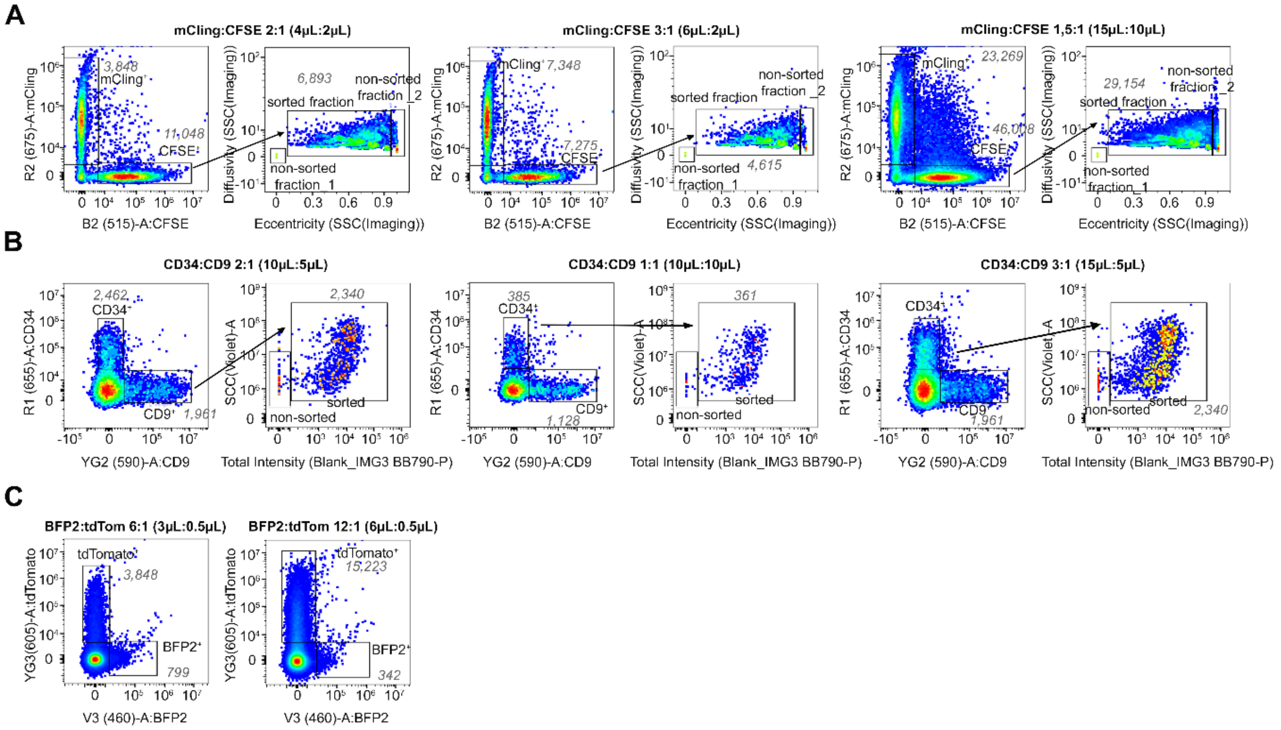


**Supplementary Figure 6. Serial dilutions of EVs within the S8 sorting gates demonstrate proportional changes in event counts, consistent with single-particle detection.** A) Serial dilutions of mCling⁺ and CFSE⁺ EVs; B) Serial dilutions of CD34⁺ and CD9⁺ EVs; C) Serial dilutions of tdTomato⁺ and BFP⁺ EVs.

### SUPPLEMENTARY FIGURE 7

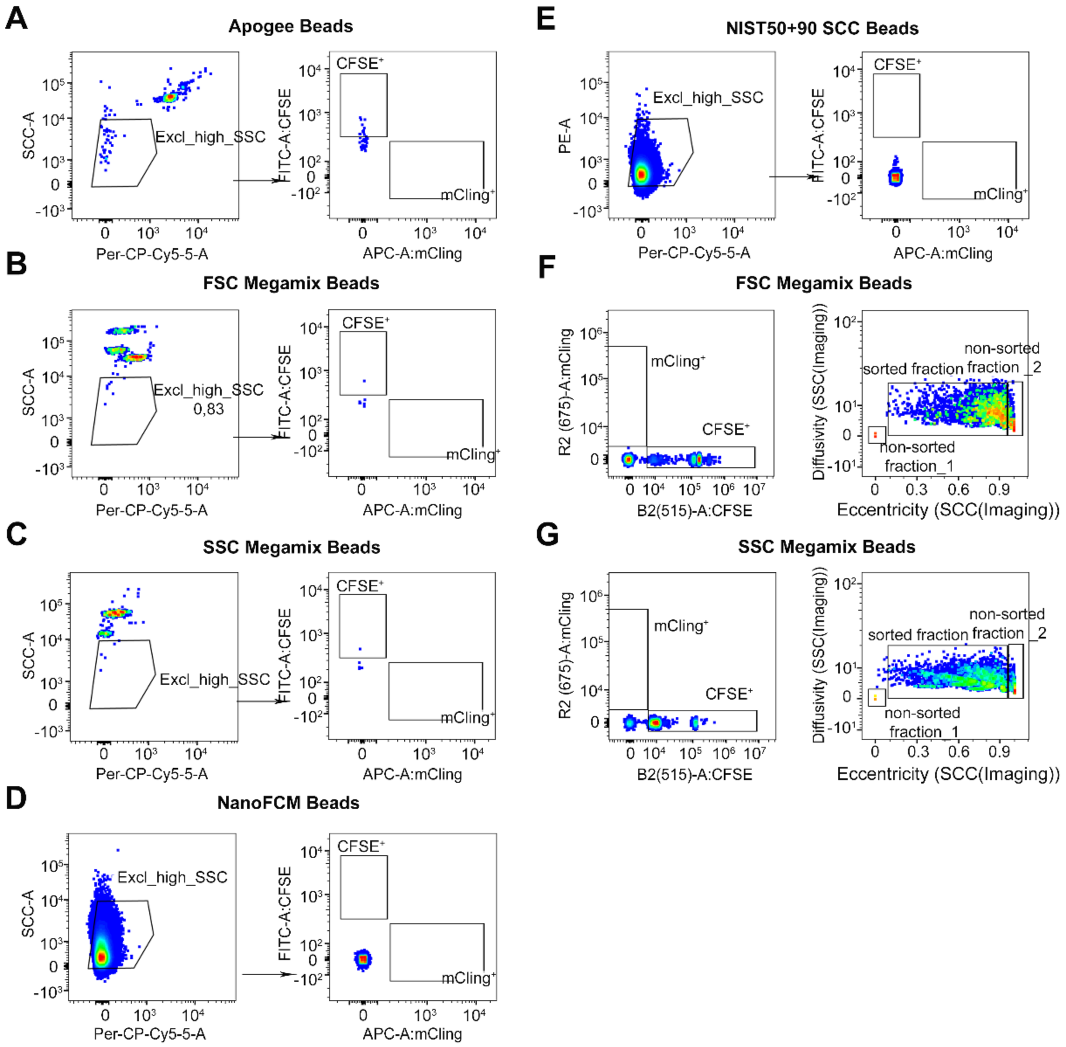


**Supplementary Figure 7. Reference beads analyzed within the sorting gates of brain tissue-derived EVs**. A) Apogee beads (starting with 80nm) within the sorting gate of the Fusion; B) FSC Megamix beads (starting with 100nm) within the sorting gate of the Fusion; C) SSC Megamix beads wihtin the sorting gate of the Fusion; D) NanoFCM beads (starting with 68nm) within the sorting gate of the Fusion; E) NIST50+90nm beads within the sorting gate of the Fusion; F) FSC Megamix Beads wihtin the sorting gate of the S8; G) SSC Megamix Beads within the sorting gate of the S8.

### SUPPLEMENTARY FIGURE 8

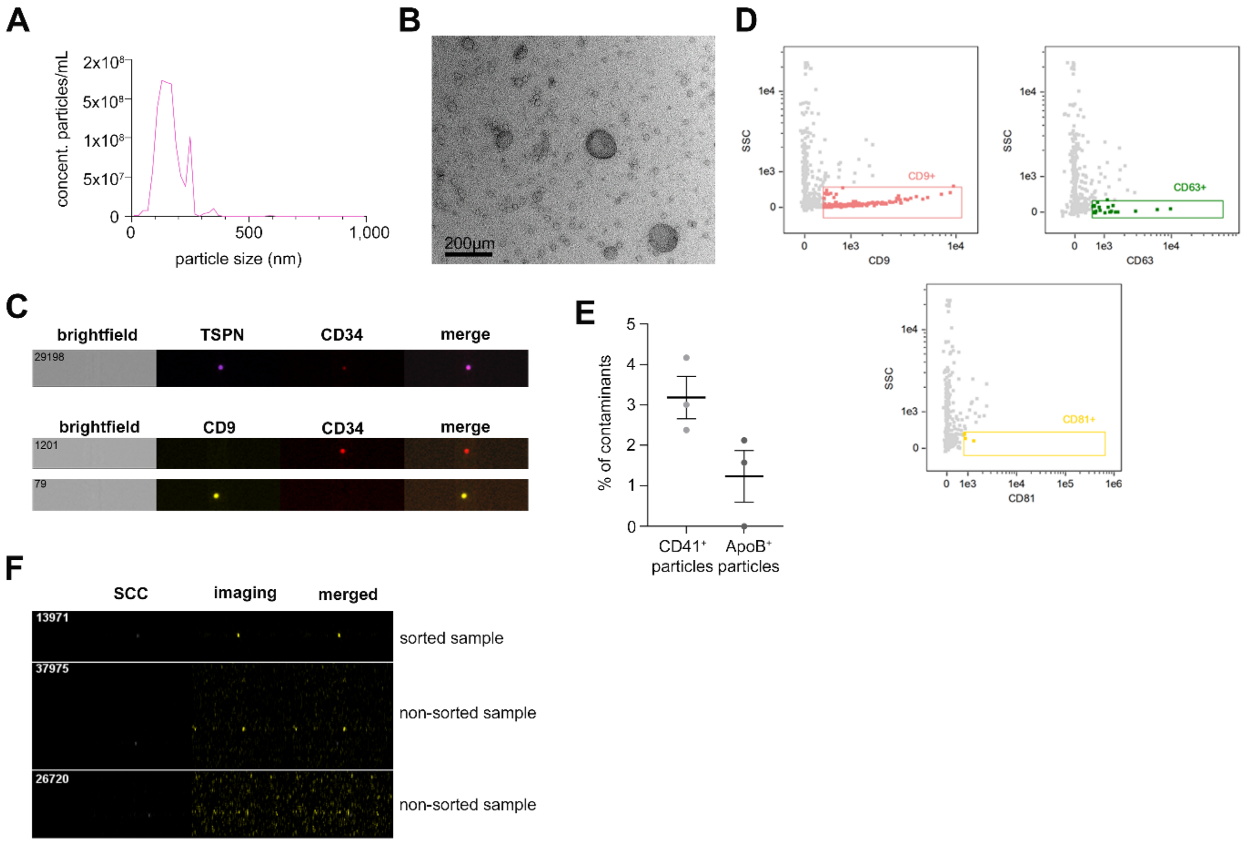


**Supplementary Figure 8. Characterization of human blood-circulating EVs**. A) NTA before sort; B) TEM of EVs before sort; C) IFCM characterization of CD34$^+$ and CD9$^+$ EVs before sort including TSPN as EV markers (top) and without TSPN (bottom); D) IFCM plots showing the expression of TSPN before sort. E) Levels of contaminants (CD41$^+$ particles and ApoB$^+$ particles) assessed via IFCM in the isolated EV fraction. Data are presented as individual points (with mean± SEM); F) Representative images of detected events within the 'sort' and 'non sort' gates of the S8, corresponding to Figure 4F.

### SUPPLEMENTARY FIGURE 9

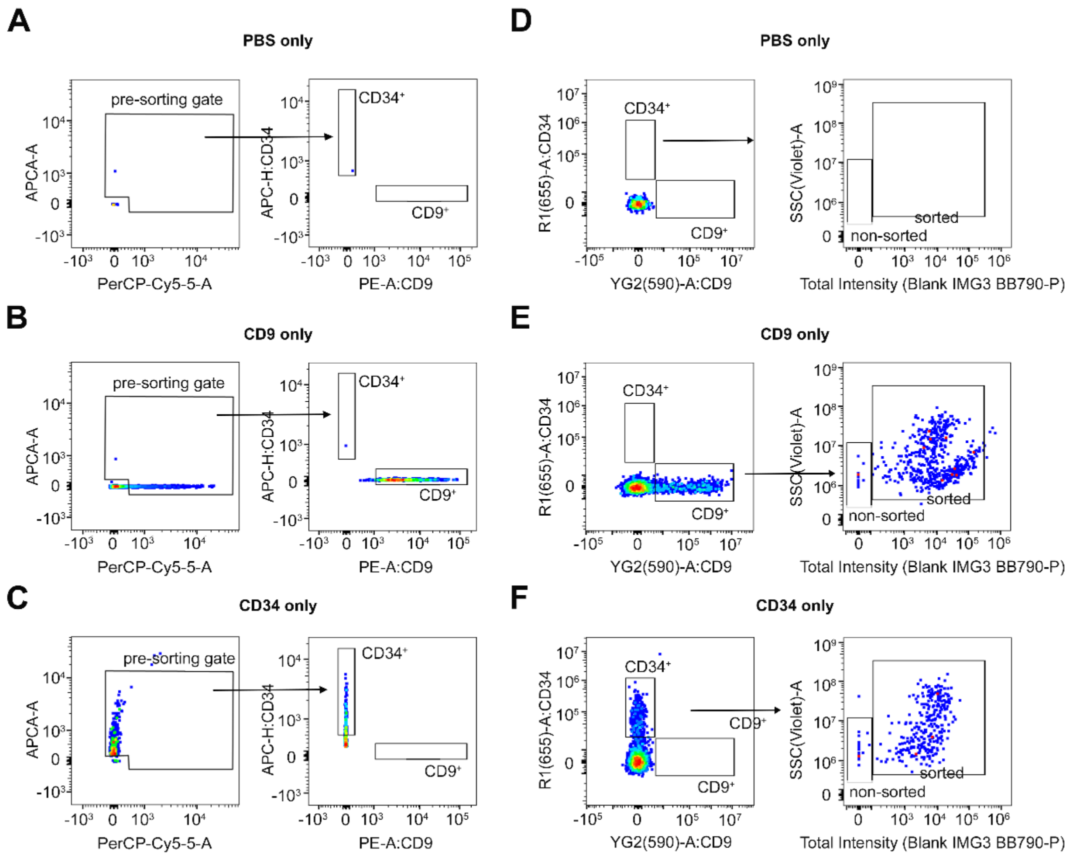


**Supplementary Figure 9. MIFlowCyt-EV sorting controls of human blood-circulating EVs (part one)**. All MIFlowCyt-EV control samples were analyzed using the same gating strategy that was applied for EV sorting. A) PBS control; B) $CD9^{+}$ (PE) EVs single control within the sorting gate of the Fusion; C) $CD34^{+}$ (APC) EVs single control within the sorting gate of the Fusion; D) PBS control within the sorting gate of the S8; E) $CD9^{+}$ EVs (PE) single control within the sorting gate of the S8; F) $CD34^{+}$ (APC) EVs single control within the sorting gate of the S8.

### SUPPLEMENTARY FIGURE 10

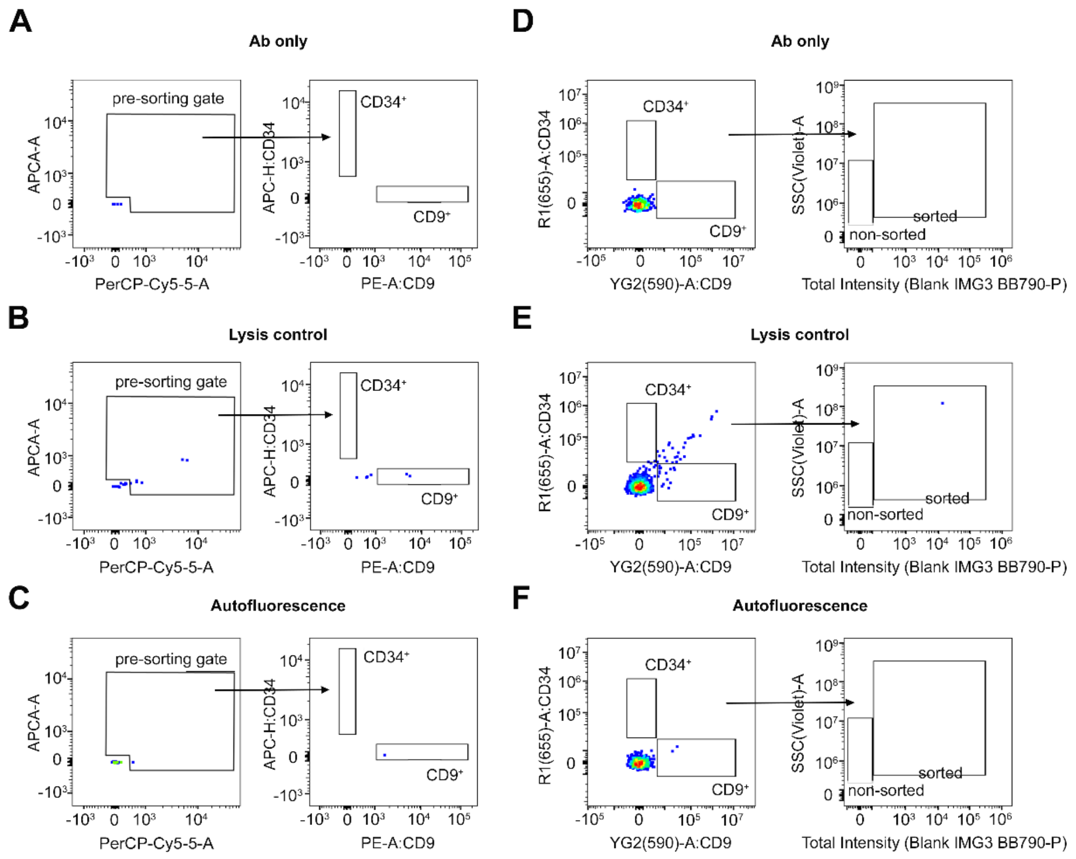


**Supplementary Figure 10. MIFlowCyt-EV sorting controls of human blood-circulating EVs (part two).** All MIFlowCyt-EV control samples were analyzed using the same gating strategy that was applied for EV sorting. A) Antibody (Ab) only control within the sorting gate of the Fusion; B) EV lysis control within the sorting gate of the Fusion; C) EVs only (autoflouresence) control within the sorting gate of the Fusion; D) Ab only control within the sorting gate of the S8; e) EV lysis control within the sorting gate of the S8. f) EVs only (autoflouresence) control within the sorting gate of the S8.

### SUPPLEMENTARY FIGURE 11

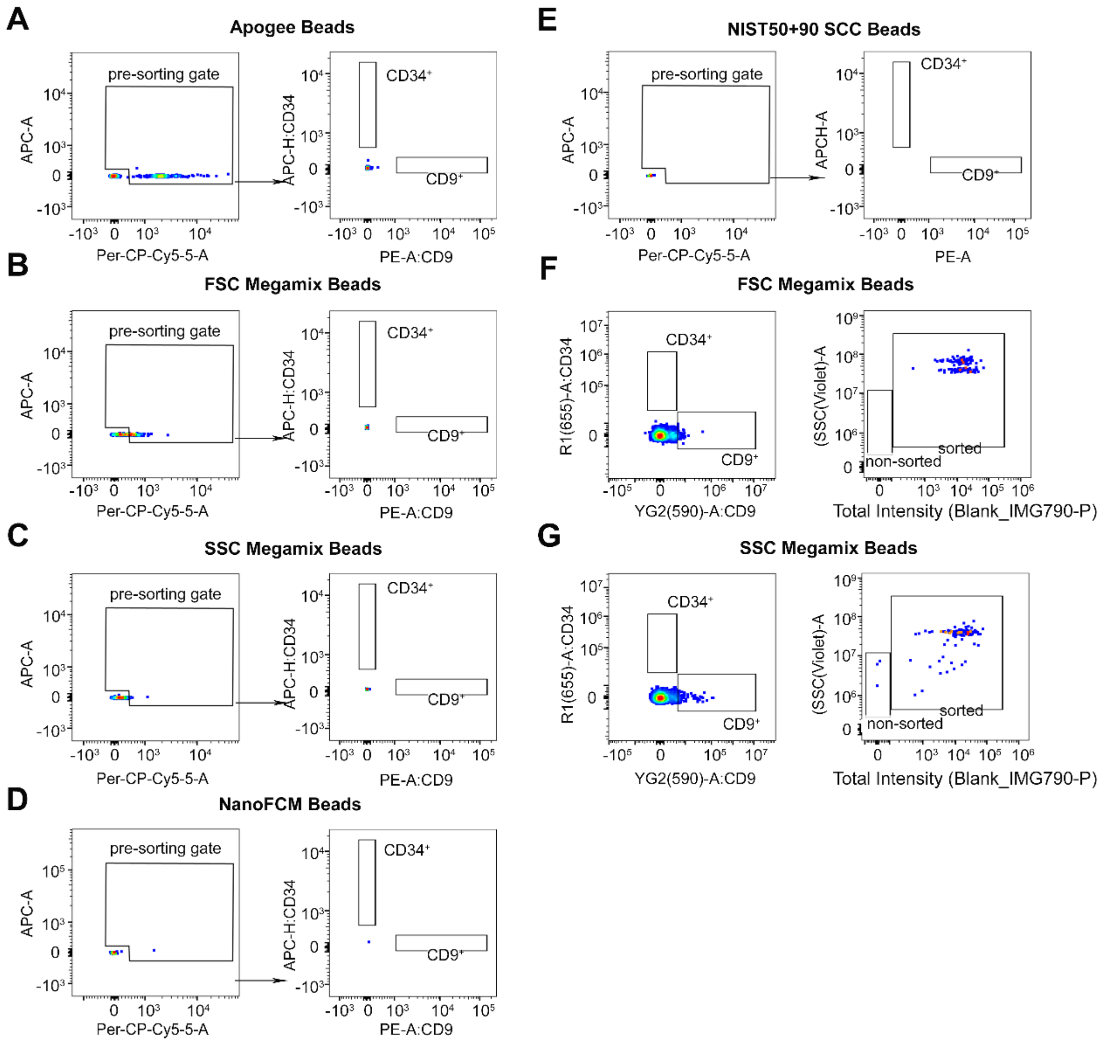


**Supplementary Figure 11. Reference beads analyzed within the sorting gates of brain tissue-derived EVs**. A) Apogee beads (starting with 80nm) within the sorting gate of the Fusion; B) FSC Megamix beads (starting with 100nm) within the sorting gate of the Fusion; C) SSC Megamix beads wihtin the sorting gate of the Fusion; D) NanoFCM beads (starting with 68nm) within the sorting gate of the Fusion; E) NIST50+90nm beads within the sorting gate of the Fusion; F) FSC Megamix Beads wihtin the sorting gate of the S8; G) SSC Megamix Beads within the sorting gate of the S8.

**SUPPLEMENTARY FIGURE 12**

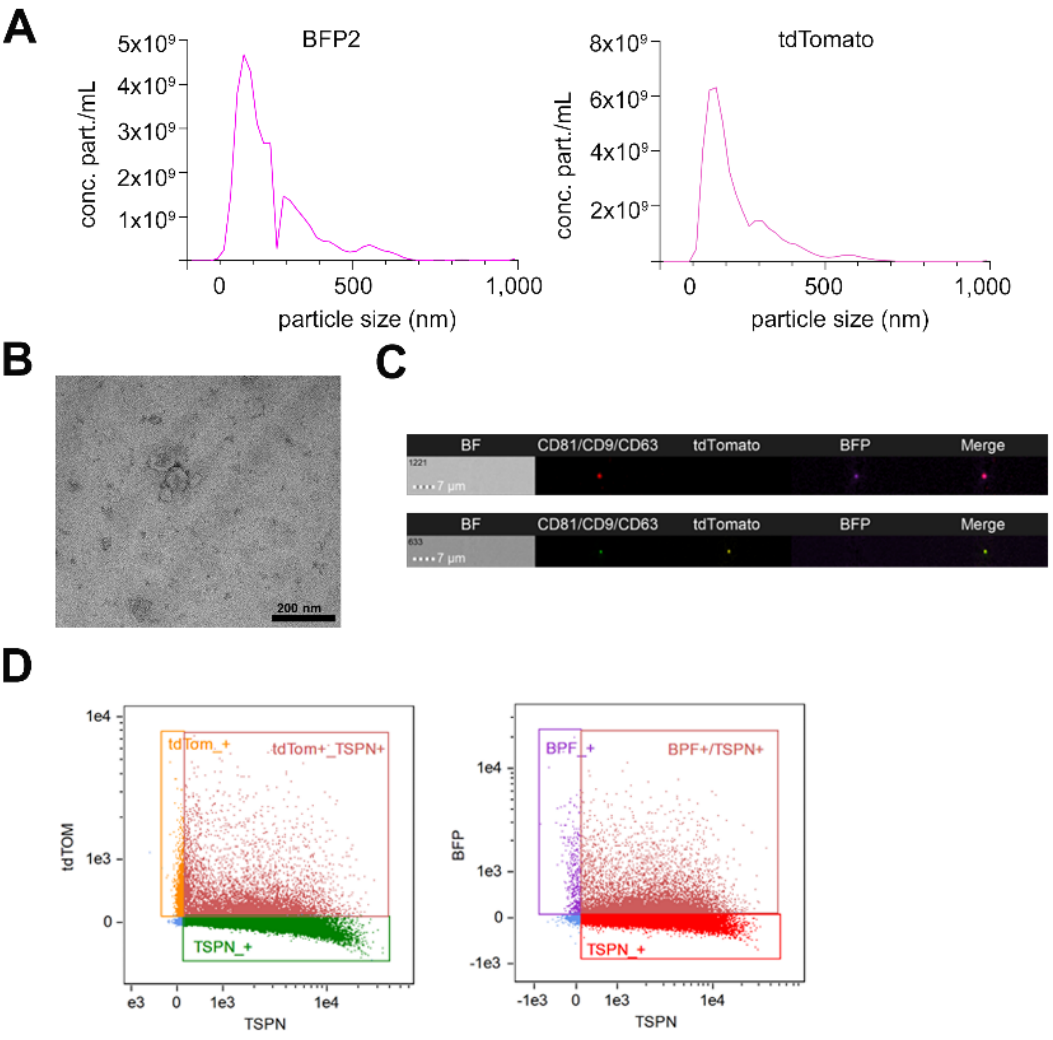


**Supplementary Figure 12. Characterization of glioma cell culture-derived EVs**. A) NTA of BFP$^{+}$ EVs (left) and tdTomato$^{+}$ EVs (right) before sort; B) TEM before sort; C) IFCM characterization of BFP$^{+}$ and tdTomato$^{+}$ EVs before sort including tetraspanins (TSPN) as EV markers (top) and without TSPN (bottom); D) IFCM plots showing the co-expression of TSPN and tdTom$^{+}$/ BFP$^{+}$ before sort.

### SUPPLEMENTARY FIGURE 13

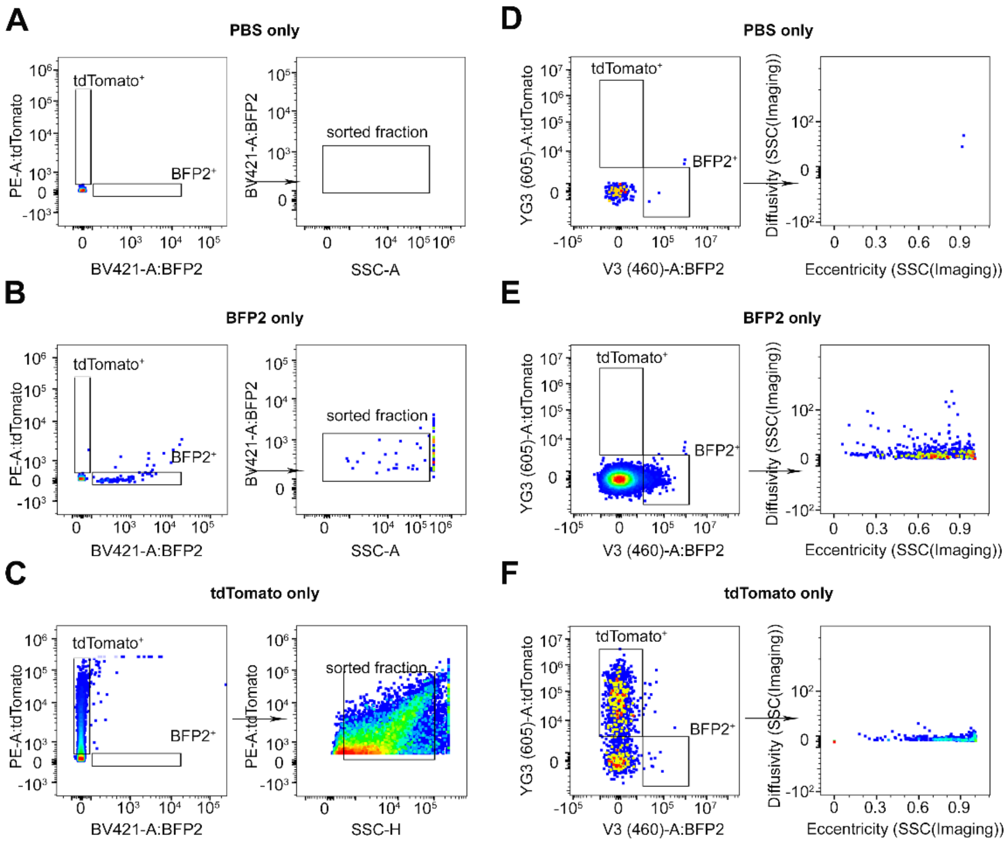


**Supplementary Figure 13. MIFlowCyt-EV sorting controls of glioblastoma cell culture-derived EVs. ).** All MIFlowCyt-EV control samples were analyzed using the same gating strategy that was applied for EV sorting; A) PBS control within the sorting gate of the Fusion; B) BFP+ EVs single control within the sorting gate of the Fusion; C) tdTomato+ EVs single control within the sorting gate of the Fusion. D) PBS control within the sorting gate of the S8; E) BFP+ EVs single control within the sorting gate of the S8; F) tdTomato+ EVs single control within the sorting gate of the S8.

### SUPPLEMENTARY FIGURE 14

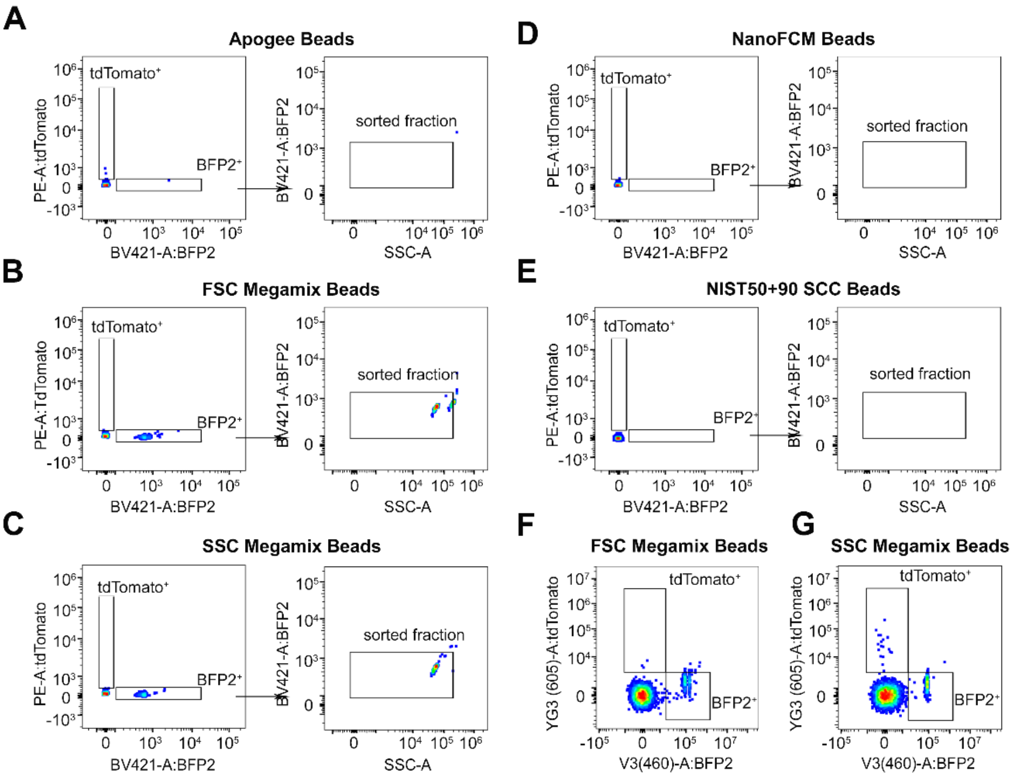


**Supplementary Figure 14. Reference beads analyzed within the sorting gates of glioblastoma cell culture-derived EVs.** A) Apogee beads (starting with 80nm) within the sorting gate of the Fusion; B) FSC Megamix beads (starting with 100nm) wihtin the sorting gate of the Fusion. C) SSC Megamix beads (starting with 160nm) within the sorting gate of the Fusion; D) NanoFCM beads (starting with 68nm) within the sorting gate of the Fusion. E) NIST 50+90nm beads within the sorting gate of the Fusion; F) FSC Megamix beads within the sorting gate of the S8; G) SSC Megamix beads within the sorting gate of the S8.

**SUPPLEMENTARY FIGURE 15**

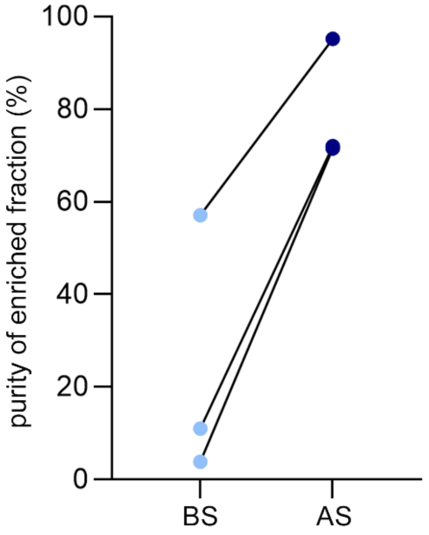


**Supplementary Figure 15. EV sort is reproducible and robust**. $CD34^+$ EVs, $BFP^+$ EVs, and $mCling^+$ EVs were independently enriched from separate experimental runs and sample batches. The percentages of each target EV population relative to the total EV population is shown before sorting (BS) and after sorting (AS), demonstrating reproducible enrichment across independent sorting experiments.

**SUPPLEMENTARY FIGURE 16**

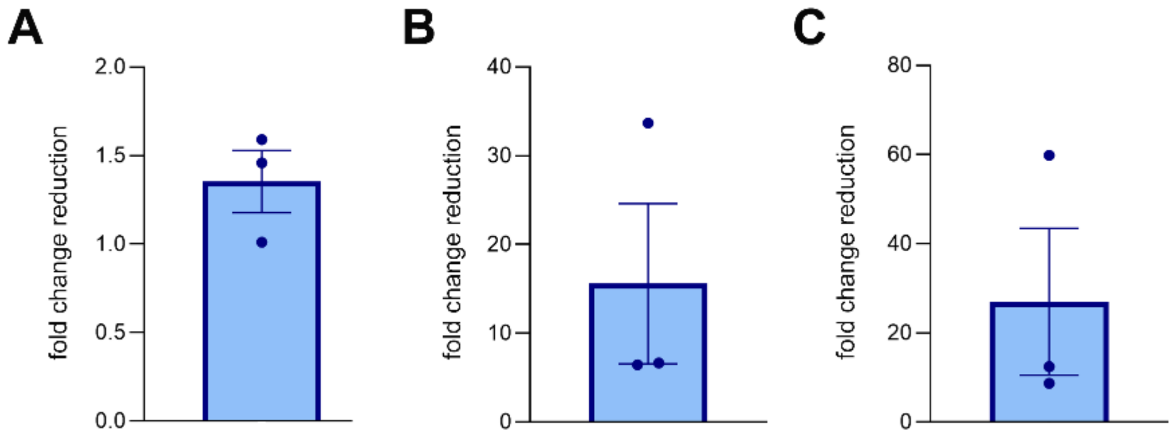


**Supplementary Figure 16. Recovery of EVs after sort.** Sorting trials with the Fusion and the S8 .All post-sort EV recoveries are expressed as fold change relative to the input EV population; A) Recovery of tissue brain-derived EVs; B) Recovery of glioma cell culture-derived EVs; C) Recovery of human blood-circulating EVs

**SUPPLEMENTARY FIGURE 17**

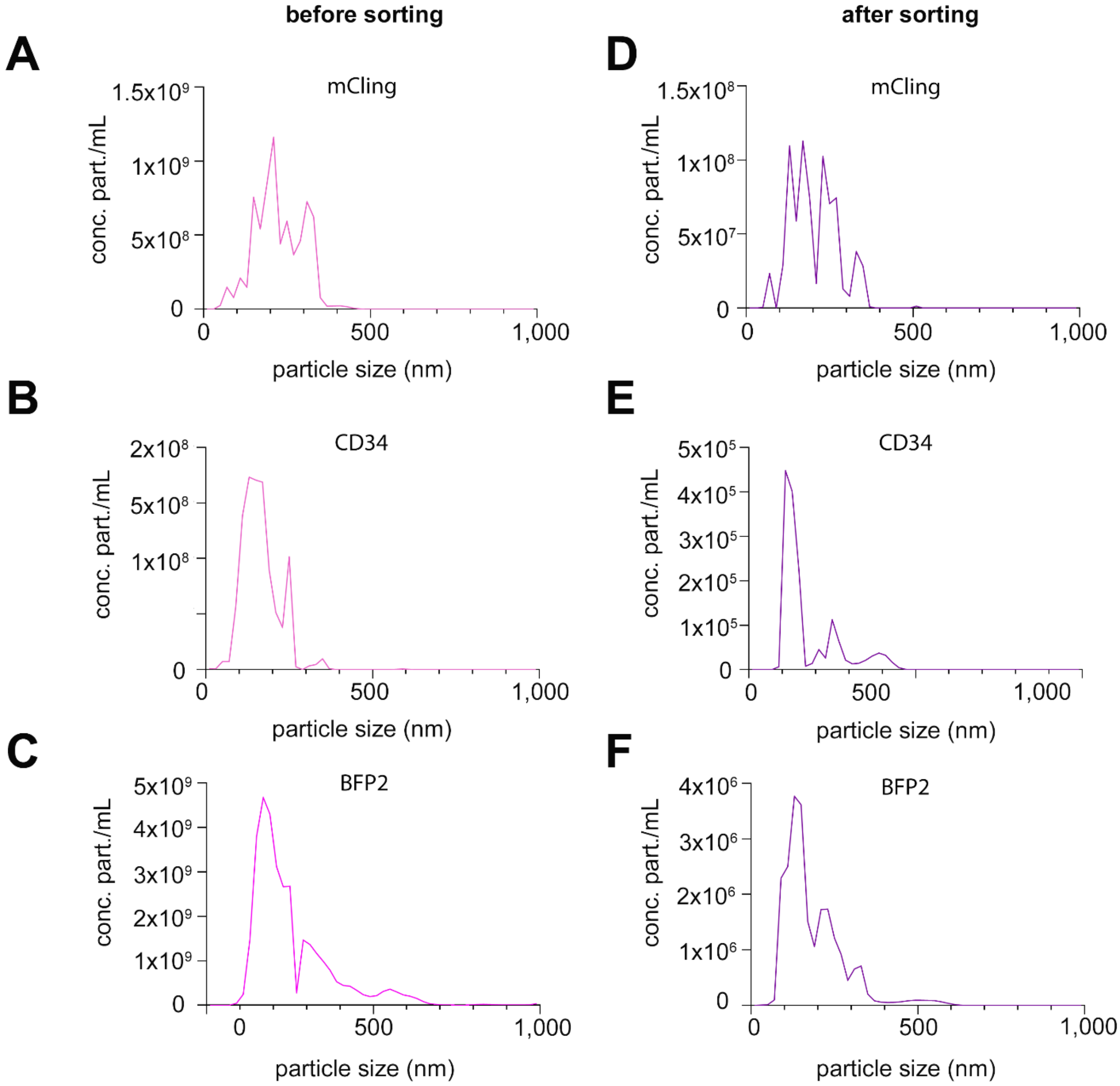


**Supplementary Figure 17. EV size is unchanged after sorting**. A) NTA measurements of mCling$^{+}$ EVs before sort; B) NTA measurements of CD34$^{+}$ EVs before sort; C) NTA measurements of BFP2$^{+}$EVs before sort. D) NTA measurements of mCling$^{+}$ EVs after sorting; E) NTA measurements of CD34$^{+}$ after sort; F) NTA measurements of BFP2$^{+}$EVs after sorting.

**SUPPLEMENTARY FIGURE 18**

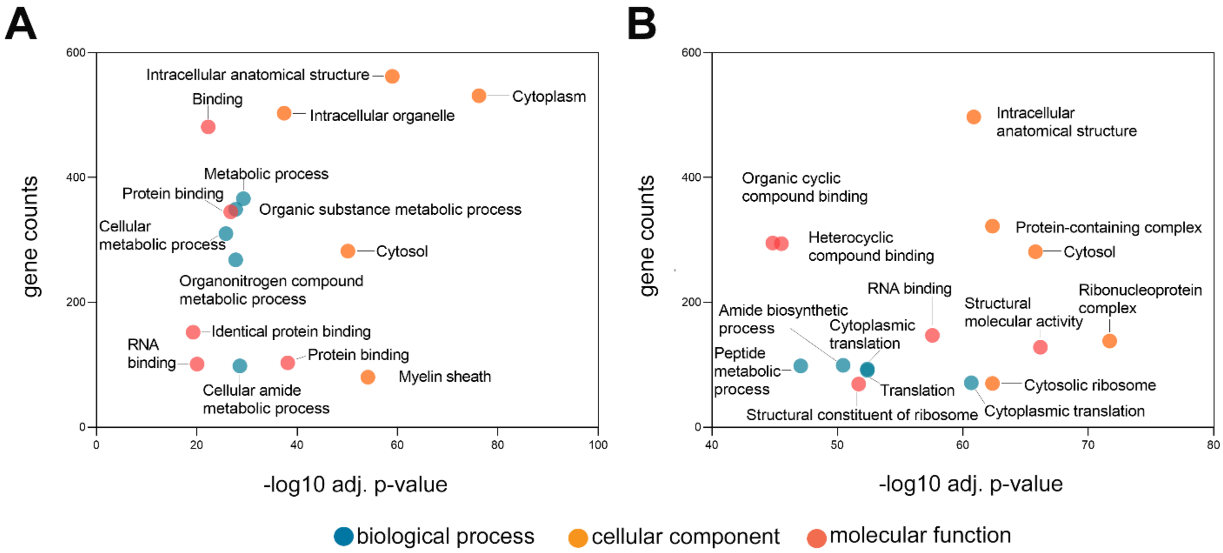


**Supplementary Figure 18. Proteomic analysis of CFSE+ EVs after sort**. A)Gene ontology analysis of CFSE$^{+}$ EVs after sort with Fusion. B) Gene ontology analysis of CFSE$^{+}$ EVs after sort with S8.
