## Supplementary material for "Enrichment of Extracellular Vesicles Subsets from Diverse Biological Sources Using Conventional and Image-Based Fluorescence Activated Sorting": MiBloodEV supplementary data: MIBloodEV_Suppl. Table 1 Graf et al JEV.pdf

INTERNATIONAL SOCIETY for EXTRACELLULAR VESICLES

### MIBlood-EV

#### Standardized Reporting Tool for Blood EV Research (Human)

#### STUDY INFORMATION

| | | | |
|---|---|---|---|
| 1.0 Manuscript title | High-Purity Enrichment of Extracellular Vesicles from diverse sources by Conventional and Image-Based Fluorescence Activated Cell Sorters for robust Downstream Applications | | |
| 1.1 Corresponding author (Name and Email) | Berta Puig | | |
| 1.2 Institution name | University Medical Centre Hamburg-Eppendorf | | |
| 1.3 Time period of experiment (e.g. 2022-2024) | 2023-2025 | 1.4 Number of samples | 30 |
| 1.5 Cargo of interest | ☑ Vesicles ☑ Protein ☐ RNA ☐ DNA ☐ Other: | | |
| 1.6 Biospecimen type | ☐ Plasma ☑ Serum | 1.7 Biospecimen state | Frozen |
| 1.8 Source of frozen specimens | Prepared by the lab | 1.9 Years of collection (range) | 2 |

#### BLOOD COLLECTION AND PROCESSING

| | | | |
|---|---|---|---|
| 2.0 Patient fasting status | Uncertain | 2.1 Fasting length (e.g. hours/days) | |
| 2.2 Anatomical access site | antecubital fossa | 2.3 Needle diameter (e.g. gauge) | 21 |
| 2.4 Blood volume collected (mL) | 4.9 | | |
| 2.5 Plasma anticoagulant | ☐ EDTA ☐ Citrate ☐ Heparin ☐ Other: | | |
| 2.6 Serum tube type | Clot activator + Gel separator (SST) | 2.7 Serum clotting time (minutes) | |
| 2.8 Time between collection and first centrifugation (range in hours) | 1-2h | | |
| 2.9 Transport temperature | Room temp (~22°C) | 2.10 Transport condition of tubes | Vertical |
| 2.11 Centrifuge brand and model | Hettich 380R centrifuge | | |
| 2.12 Bucket rotor type | Fixed | 2.13 Number of centrifugation cycles | One |
| 2.14a 1st Centrifugation speed (RCF in x g) | 2000 | 2.14b 1st Centrifugation time (minutes) | 15min |
| 2.15 1st Rotor brake | Off | 2.16 1st Centrifugation temperature | Room temp (~22°C) |
| 2.17a 2nd Centrifugation speed (RCF in x g) | | 2.17b 2nd Centrifugation time (minutes) | |
| 2.18 2nd Rotor brake | | 2.19 2nd Centrifugation temperature | |
| 2.20 Additional processing steps (e.g. filtration) | | | |
| 2.21 Storage tubes (brand, type, source, catalog number) | Eppendorf Tubes 1.5ml | | |
| 2.22 Storage temperature | -80°C | 2.23 Length of storage (range in years) | max. 2 years |

#### PLASMA/SERUM QUALITY CONTROL

| | | | |
|---|---|---|---|
| 3.0 Number of freeze-thaw cycles (range) | 0 | | |
| 3.1 Thawing temperature | on ice | 3.2 Thawing duration (minutes) | over night |

#### Hemolysis

| | | |
|---|---|---|
| 3.3 Presence of hemolysis | No | 3.4 Frequency of hemolyzed samples (e.g. <25%, 25-50%) |
| 3.5 Method used | Visual inspection | 3.6 RBC count (Median, 95% CI, N) |
| 3.7 RBC counter brand and type | | |
| 3.8 Spectrophotometry hemoglobin concentration (mean g/L) | | |
| 3.9 Spectrophotometer brand, model and wavelength measured (e.g. 414 nm) | | |
| 3.10 Hemolized samples were discarded | Yes | |

#### Platelets

| | | | |
|---|---|---|---|
| 3.11 **Presence of platelets** | Yes | 3.12 **Method used (e.g. Flow Cytometry)** | Flow Cytometry |
| 3.13 **Marker(s) used (e.g. CD61, CD41)** | CD41 | | |
| 3.14 **Concentration (median, 95% CI, N)** | 3.0% of CD9+ particles (95 CI 2.38;4.17; n=3) | | |
| 3.15 **Platelet counter instrument brand, type and limit of detection (cells/L)** | | | |
| 3.16 **Flow cytometer brand and type** | Cytek Amnis ImageStream Mk II | | |
| 3.17 **Flow cytometry size and fluorescence ranges of detection in nanometers and MESF, respectively** | | | |

#### Lipoproteins

| | | | |
|---|---|---|---|
| 3.18 **Presence of lipoproteins** | Yes | 3.19 **Method used (WB, ELISA, FC)** | Flow cytometry |
| 3.20 **Spectrophotometry L-index** | | | |
| 3.21 **Spectrophotometer brand, model and wavelength measured (e.g. 700 nm)** | | | |
| 3.22 **WB Marker(s) used (e.g. Apo B)** | | | |
| 3.23 **Western blot images provided in manuscript?** | | | |
| 3.24 **Flow cytometry marker(s) used (e.g. ApoB)** | ApoB | | |
| 3.25 **Flow cytometry concentration (median, 95% CI, N)** | 1.57% of CD9+ particles (95 CI 0.00;2.13; n=3) | | |
| 3.26 **Flow cytometer brand and type** | Cytek Amnis ImageStream Mk II | | |
| 3.27 **Flow cytometry size and fluorescence ranges of detection in nanometers and MESF, respectively** | | | |
| 3.28 **Brand and catalog number for ELISA kit used for each Apolipoprotein test (e.g. ApoB)** | | | |
| 3.29 **Concentration measured for each Apolipoprotein tested (mean µg/mL)** | | | |
